# Tracking long-term functional connectivity maps in human stem-cell-derived neuronal networks by holographic-optogenetic stimulation

**DOI:** 10.1101/2021.05.11.443589

**Authors:** Felix Schmieder, Rouhollah Habibey, Johannes Striebel, Lars Büttner, Jürgen Czarske, Volker Busskamp

**Affiliations:** Laboratory of Measurement and Sensor System Technique, Faculty of Electrical and Computer Engineering, TU Dresden, Helmholtzstraße 18, 01069 Dresden, Germany; Universitäts-Augenklinik Bonn, University of Bonn, Dep. of Ophthalmology, Ernst-Abbe-Straße 2, D-53127 Bonn, Germany; Competence Center for Biomedical Computational Laser Systems (BIOLAS), TU Dresden, Dresden, Germany; Cluster of Excellence Physics of Life, TU Dresden, Dresden, Germany; Institute of Applied Physics, School of Science, TU Dresden, Dresden, Germany

**Keywords:** Functional connectivity maps, Computational holographic stimulation, Human stem-cell-derived neurons, Long-term neuronal network culture, Multi-electrode array electrophysiology, Optogenetics.

## Abstract

Neuronal networks derived from human induced pluripotent stem cells (hiPSCs) have been exploited widely for modelling neuronal circuits, neurological diseases and drug screening. As these networks require extended culturing periods to functionally mature *in vitro*, most studies are based on immature networks. To obtain insights on long-term functional features of human networks, we improved a long-term glia-co-culture culturing protocol directly on multi-electrode arrays (MEA), facilitating long-term assessment of electrical features at weekly intervals. We applied optogenetic stimulation to induce neuronal activity, which resulted in accelerated neuronal responses during network development. Using holographic stimulation with single-cell-resolution, propagating evoked activities of 400 individually stimulated neurons per MEA were traceable, and precise network functional connectivity motifs were revealed. Our integrated holographic optogenetic stimulation platform on MEAs facilitates studying long-term functional dynamics of human neuronal networks *in vitro*. This is an important step towards establishing hiPSC-derived neurons as profound functional testbeds for basic and biomedical research.

**Highlights:**

1. Integrated platform allowed long-term optogenetic experiments on hiPSC-derived networks.
2. Full-field optogenetic stimulation boosted hiPSC-derived neuronal network activity.
3. Single-neuron resolution holographic stimulation evoked local responses in the network.
4. Holographic stimulation of each neuron revealed its functional connectivity patterns.
5. Subsequent holographic stimulation of more than 400 neurons revealed the whole network connectivity map.

## Introduction

Complex neuronal circuits of the human brain are composed of diverse neuronal cell types and their elaborate connections (1, 2). The functional connectivity patterns of the developing and adult human brain are a complex system of interacting units (1, 3). Determining how a single neuron contributes to circuit function and behavior is a key step towards unraveling functional connectivity maps within neuronal circuits (4, 5). Animal models have been utilized extensively to explore neuronal networks, and have provided substantial understanding of their functional and structural organization (6, 7). However, the emerging field of human pluripotent stem cell (hPSC) technology enables bottom-up neuroscientific approaches by generating neurons and neuronal networks *in vitro*. Especially for biomedical studies, this alternative route is complementary to animal model systems and has the potential to reduce the translational gap from bench-to-bedside (8). Human embryonic (hESCs) (9) and human induced pluripotent stem cells (hiPSCs) (10, 11) offer an almost unlimited cell source for neuronal cell and circuit engineering. There are many methods available to drive stem cells into neurons of interest. Many protocols require multiple steps and long time periods using soluble factors and specific culturing techniques (12), but single-step transformations by inducible neurogenic transcription factor expression can result in rapid neurogenesis (13–15). For instance, there are transgenic hiPSCs in which neurogenin-1 and neurogenin-2 can be induced by the TetOn promoter system, so called iNGN cells: these have been extensively characterized. These cells are highly proliferative as hiPSCs and the neurogenic transcription factor activation triggers neurogenesis, leading to postmitotic neurons in just 4 days post induction (dpi). This process is independent of the underlying stem cell line used (13).

Because *in-vivo* maturation of neuronal circuits occurs naturally, sophisticated functional studies are feasible and have shed light on many brain processing features (16). However, to date, the electrophysiological properties of most human stem-cell-derived neurons are insufficiently characterized. Most *in-vitro* cultures mature slowly (17, 18). Therefore, neuronal activity is frequently induced by current injection at a few time points to study some functional aspects (18). Continuous recordings of the developing human neuronal circuits are rare. One reason for this is the challenging nature of patch-clamp recordings: sterility cannot be guaranteed, requiring the disposal of the cultures after one experiment (19). Multielectrode arrays (MEAs) allow for recording from identical cultures over time through their non-invasive and sterile working mode (20, 21). MEA technology has enabled researchers to continuously study the network properties of developing primary cortical and hippocampal neuronal cultures (22, 23) and hiPSC-derived neuronal networks (24). Long-term activity profiles of the commercially available hiPSC-derived “iCell” neurons were studied using standard MEAs (25, 26) and high-density CMOS-based MEAs (18). These studies have revealed increased neuronal firing rates, i.e. action potential (AP) frequencies, over time (25), while the earliest detection of synchronized burst activity with higher latencies was at day 70 of culture (18, 27).

Long-term functional properties of single iNGN neurons have been characterized using whole-cell patch-clamp electrophysiology recordings at different dpi: these cells respond to current injection from 14 dpi onwards, and develop excitatory glutamatergic synapses over time. However, robust spontaneous activity first developed at 56 dpi (28). Another functional study in which the iNGN cassette was introduced into hESCs detected an increasing trend of neuronal network firing rates and synaptic transmission over two months using patch-clamp and MEA electrophysiology (25). In addition, a small fraction of neurons developed into inhibitory GABAergic neurons. As functional maturation of hiPSC-derived neurons in general, and in particular for iNGN cells, takes a long time, neuronal culturing techniques have been improved to promote long-term neuronal survival. Astrocyte co-cultures (12, 29) or astrocyte-conditioned media (28) have been shown to be essential to support the fragile human neurons *in vitro* in long-term cultures. Besides recording spontaneous network activity, functional responses can also be invoked using MEA stimuli (18). MEA recording electrodes are then used to apply voltage or current pulses. The major drawbacks of this method are the low spatial resolution of the applied stimuli in the network, and stimulation artifacts (30). For example, no meaningful activity data can be extracted while the stimulation pulse is being applied, nor for a few milliseconds afterwards, (30). Here, optogenetics represents a valuable tool to provide precise spatial resolution of the stimulation and to avoid stimulation artifacts (31).

Optogenetics refers to DNA-encoded light-sensitive ion channels or pumps, which use light stimuli to facilitate the control of neuronal activity when ectopically expressed in neurons (32). Channelrhodopsin-2 (ChR-2) is among the most commonly-used optogenetic actuators for controlling neuronal cell activity by light-induced depolarization (33, 34). Optogenetic actuators are usually tagged with a fluorescent protein such as enhanced yellow fluorescent protein (EYFP) to visualize the expression within the membrane of targeted cells (35). The major advantage of optogenetic stimulation is the possibility to express different types of optogenetic actuators in specific neuronal populations (36–38), and to apply focused or patterned light stimuli, which requires advanced stimulation and regulation technology, to target specific neurons within a defined neuronal network (39). Stimulation of individual neurons, and tracking the resulting activity propagation across the neuronal networks, makes it easier to study network connectivity and dynamics. Advanced optogenetic measurements use two-photon stimulation to penetrate deep into the neuronal tissues of interest, not only *ex vivo* (40) but also in freely moving animals (41). Unfortunately, these two-photon stimulation setups are unaffordable for many labs. With single-photon stimulation methods, such as galvanometric actuator or acousto-optic deflector scanning techniques, it is not possible to simultaneously stimulate several neurons (42, 43). However, several parallel approaches have been reported. Arrays of microscopic organic and non-organic light-emitting diodes (LED) have been used for optogenetics (44, 45) and, in the future, these devices may shrink to the size of single-neuron soma. Due to the beam characteristics of such small emitters, without additional optics they will probably be limited to two-dimensional applications unless accompanied with structures aiding in deep-tissue stimulation (46). Moreover, the heat generated by the LEDs might harm the surrounding tissue if they are directly integrated into needle probes (47). More flexible methods use pixelated light modulation devices with millions of individually addressable actuators like digital mirror devices (DMDs) or spatial light modulators (SLMs). These devices can be used to modulate either the amplitude or phase of the light to create stimulation patterns on the sample. When the light amplitude is modulated, the highest switching rates of DMDs are several kHz, but they have a poor light efficiency because of the amplitude modulation (48, 49). When the phase of the light wave is modulated, generalized phase contrast (GPC) offers light-efficient speckle-free illumination. Unless additional techniques with increased instrumentation and adjustment effort are added, GPC is limited to two-dimensional projection (50). Single-photon stimulation with computer-generated phase-modulating holograms is more flexible than the methods mentioned above: higher stimulation energies than DMD or organic light emitting diode (OLED) illumination are possible, as well as three-dimensional scanning (48), fiber-optic approaches are easier (51, 52), and even sample-induced aberrations can be compensated for (53). As the most flexible and economical path for our experiments, we chose to use single-photon stimulation with computer-generated holograms based on a ferroelectric SLM with switching times of up to 400 Hz, providing high spatial resolution stimuli at neuronal soma size (8 µm) with sufficient temporal resolution (2.5 ms) and high stimulation energies (0.15 W/mm^2^) (54). Optogenetics has already been widely exploited for mapping microcircuits by targeting subsets of neurons in brain slices *in vitro* (55). Optogenetic-based probing of the *in vivo* inter-regional and global connections within rodent brain areas is also commonly used (56, 57). Large areas of rat primary cortical networks have also been functionally studied, in combination with optogenetic manipulation, to extract functional connectivity maps *in vitro* (58). However, optogenetic-based mapping of functional connectivity has not been explored in human stem-cell-derived neuronal networks *in vitro*. As most functional studies on hiPSC-derived neurons just provide snapshot information on action potential (AP) frequencies, in-depth information about *in vitro* neuronal network development remains elusive (28, 59).

Here we implemented a holographic stimulation device for MEA recordings to activate individual hiPSC-derived neurons and extract precise connectivity data from developing neuronal networks over three months of weekly recordings. To this end, we optimized our human neuronal culturing protocol, facilitating stable holographic stimulation experiments over hours and long-term survival. The holographic stimulation of single optogenetically-tagged neurons, on average 400 per MEA culture, evoked robust neuronal activity. This cellular one-by-one stimulation, in contrast to commonly used full-field light stimulation, facilitated the extraction of dynamic individual connectivity motifs. Our single-cell activation data provided functional insights into hiPSC-derived neuronal networks, including the extraction of single-neuron connectivity patterns, total neuronal connection numbers, and synaptic strengths. In parallel, we also applied full-field optogenetic stimulation, which led to stronger neuronal activity by driving hundreds of neurons at once. Still, our full-field stimulation data revealed that robust neuronal activity can be evoked at earlier developmental time points than the emergence of spontaneous activity. Combining single-cell optogenetics with long-term human stem-cell-derived neuronal networks within MEAs enables parallel collection of morphological and functional data. This is key to implementing these *in vitro* model systems in the field of systems neuroscience for both basic and also biomedical research.

## Results

### Stable co-culture platform for long-term optogenetic experiments on human stem-cell-derived neurons

In order to continuously record neuronal activity from iNGN neurons over months, we have established an advanced culturing protocol on MEAs (Figure 1A-C, see methods). Upon neuronal induction, iNGN cells were transduced with ChR2-EYFP lentiviral particles. The EYFP tag resulted in membrane-bound labeling of targeted neurons, making it easier to capture neuronal morphology by fluorescent live cell imaging within the MEA setup. The antimetabolic agent cytosine arabinoside (ARA-C) was added at 4 days post induction (dpi) to remove undifferentiated hiPSC cells that would otherwise grow over the postmitotic neurons over time. At 5 dpi, the differentiated iNGN cells were enzymatically detached from the culturing plate and reseeded over the electrode area in MEA chambers (Figure 1A and C). Astrocytes were added upside-down on coverslips, separated by paraffin spacers from the neurons (Figure 1B). These so-called Banker neuron-glia co-cultures (60), in combination with neuronal media and sterile membranous lids, enabled long-term cultivation and continuous MEA recordings. For each recording, an MEA culture was placed into the MEA setup. For every recording session, we performed 3 min baseline, 5 min of full-field and a 1 h holographic light stimulation recording (Figure 1A). While we found that CO_2_ levels did not need to be adjusted within the MEA setup, the temperature was set at 37°C, which was crucial to keep the neuronal cultures healthy and to repeatedly perform recordings over months. All full-field and holographic optogenetic experiments were performed with MEAs that were sealed with transparent caps to prevent media evaporation and contaminations, while enabling fluorescent imaging and optical manipulations (Figure 1B, Figure S1). Altogether, our optimized protocol facilitated continuous functional recordings covering the maturation and neuronal circuit development periods of hiPSC-derived neurons.

**Figure 1.**
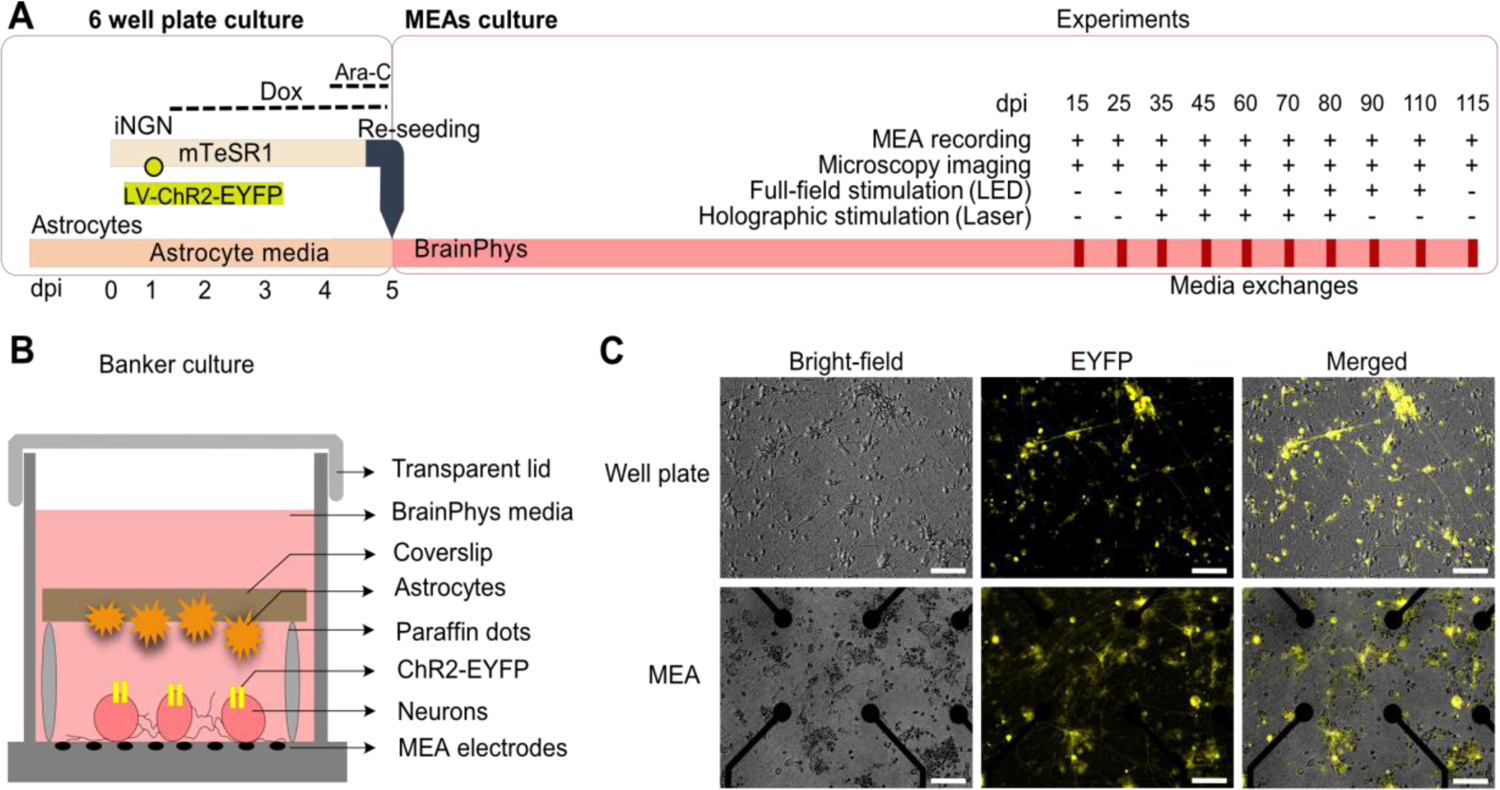
iNGN-astrocyte co-culture system on transparent MEAs. (A) Experimental scheme of iNGN-astrocyte co-culture preparation (left) and experimental procedure (right) over time. Functional recordings, microscopy imaging, full-field and holographic stimulations were studied on different days post induction (dpi). (B) Schematic of the so-called Banker culture, a neuron-glia co-culture system within the MEA device. (C) Representative microscopy images of iNGN cells. Upper panel, iNGN cells grown on Matrigel-coated well plates at 3 dpi. Lower panel, PDL-laminin-coated MEA chambers at 15 dpi. ChR2-EYFP-labeled cells are visualized by live cell microscopy. Scale bars, 100 µm. mTeSR1: standardized medium for the feeder-independent maintenance of hiPSCs. BrainPhys: serum-free neurophysiological basal medium for improved neuronal function. LV-ChR2-EYFP: lentiviral particles delivering ChR2-EYFP to iNGN cells. Dox: doxycycline. Ara-C: cytosine arabinoside to remove undifferentiated cells.

### Neuronal activity evoked by full-field optogenetic stimulation

Spontaneous neuronal activity of hiPSC-derived neurons is generally considered to be a sign of neuronal maturation, which can take weeks to months *in vitro*. Indeed, we detected signs of spontaneous iNGN activity at 25 dpi; this activity increased over time (Figure 2A). To overcome the well-known issues of electrical stimulation, we used optogenetics to replace electrical with light stimulation. Long-term spontaneous and optically-evoked activity was collected (N=4 MEAs and n=95 active electrodes, from 15 to 80 dpi). Full-field light stimulation (470 nm, 50 ms pulses, 0.5 Hz, 0.46 mW/mm^2^; Figure S1) induced APs starting at 15 dpi when unstimulated neurons were inactive and not showing signs of spontaneous activity (Figure 2D). As revealed by spike sorting, individual electrodes captured the activity of several neurons (Figure 2C). The AP wavefronts were similar between spontaneous and light-induced activity. Consistently, more neurons were active when full-field light stimulation was applied (Figure 2C, Figure S2 and Figure S3).

**Figure 2.**
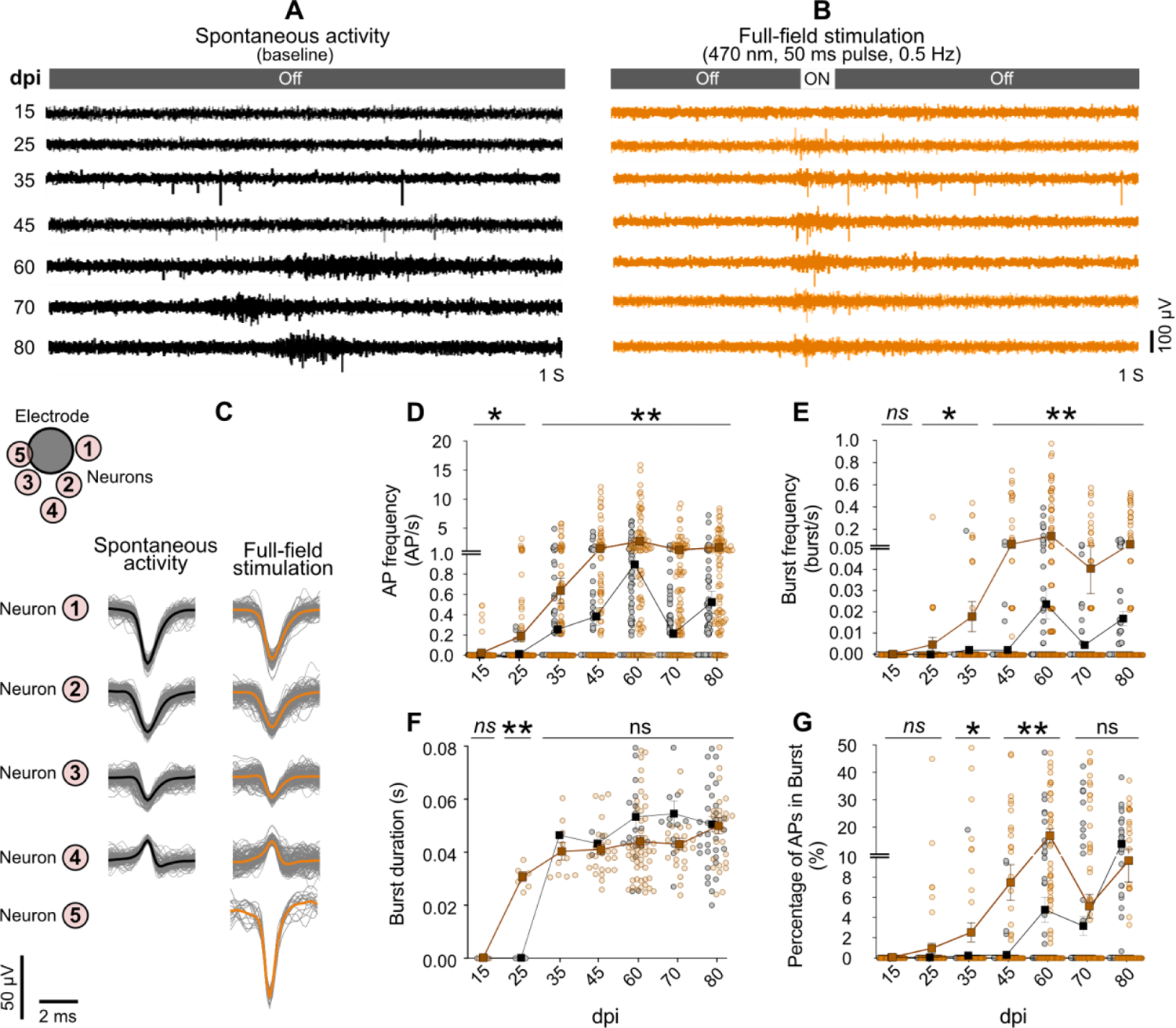
Full-field optogenetic stimulation elevated network activity features (orange) compared to spontaneous activity (black). (A) Spontaneous activity from a representative electrode over time, from 15 to 80 dpi. (B) Full-field stimulation: optically evoked activity in the same electrode on different days. (C) Action potentials of different neurons measured by one electrode were extracted from the recorded baseline and full-field stimulation signals by spike sorting. Each waveform represents activity recorded from one neuron (1–4). Neuron number 5 was only active during full-field stimulation. (D-G) The effect of full-field optogenetic stimulation on network functional features over time. All full-field stimulation data presented here was averaged over stimulation and non-stimulation time intervals and across 4 MEAs (n=95 electrodes) on each individual day. Each circle represents one electrode and each square the average for that day. Action potential frequency (D), burst frequency (E), burst duration (F), and percentage of action potentials that occurred during the burst (G) are shown. All data are represented as mean ± SEM. In all cases the average response to the full-field stimulation is compared with baseline activity on the same day (*p<0.05, **p<0.01, ***p<0.001, ns not significant) by Wilcoxon matched-pairs signed rank test.

Spontaneous APs started to be detectable at 25 dpi and reached an average frequency peak of 0.89±0.14 Hz at 60 dpi. At later time points, the AP frequency dropped again to 0.2±0.04 Hz at 70 dpi and 0.51±0.10 Hz at 80 dpi (Figure 2D). Developing neuronal networks typically have a peak in activity followed by a decline (61), suggesting that human stem-cell-derived neuronal networks replicate this developmental feature *in vitro*. By applying full-field optogenetic stimulation, we observed neuronal activity at earlier time points, starting at 15 dpi (p<0.05 vs. baseline, Figure 2D). Between 35 and 80 dpi, full-field stimulation significantly boosted network activity compared to spontaneous activity (p<0.01 vs. baseline Figure 2D). The average AP frequency of light-ON and light-OFF intervals reached 2.69±0.35 Hz at 60 dpi (Figure 2D). Next, we studied the appearance of AP bursts which are important signs of neuronal network maturation (62). Spontaneous burst activity was only captured by a few electrodes starting at 35 dpi and reached its peak at 60 dpi. Light-evoked burst activity was already detected at 25 dpi (p<0.05 vs. baseline), increased over time, and reached its peak at 60 dpi (p<0.001 vs. 15 and 25 dpi, and p<0.01 vs. baseline; Figure 2E). The average burst duration remained stable over time and was no different between spontaneous and optogenetic-evoked activity (Figure 2F). The percentage of APs appearing in bursts rather than individually increased with culture age (Figure 2G).

In order to distinguish between directly stimulated neurons and responses resulting from poly-synaptic excitation, we applied pharmaceutical inhibition. We added the excitatory glutamatergic synaptic blockers NBQX (2,3-dioxo-6-nitro-7-sulfamoyl-benzo[f]quinoxaline) and APV ((2R)-amino-5-phosphonovaleric acid) (Figure S4) to the cultures, which resulted in a decrease in APs and burst frequencies (p<0.001 vs. baseline; Figure S4). Full-field optogenetic stimulation increased AP frequency only in a few electrodes. Even higher intensities of full-field stimulation were not able to return synchronized burst activity in the presence of NBQX+APV (p<0.001 vs. baseline in untreated conditions; Figure S4). After washout, AP frequency dramatically increased under both baseline and light-stimulated conditions (p<0.05 vs. before treatment values; Figure S4B). Burst activity also appeared again after washout under light-stimulated and baseline conditions (Figure S4C).

Our long-term MEA electrophysiology data revealed different functional features of developing iNGN networks. Neuronal activity shifted from local and sparse APs to synchronized burst activities over time (Figure 2A and Figure 2E). Optogenetic stimulation also activated neurons that were silent when capturing spontaneous activity. In addition, optogenetic stimulation increased the iNGN firing rates of active neurons at earlier developmental time points. Hence, full-field optogenetic stimulation is very beneficial for boosting the detection of neuronal activity of hiPSC-derived neurons over development.

### Precise optogenetic activation of single neurons by holographic stimulation

While full-field optogenetic stimulation resulted in robust neuronal network responses, precise sub-network activities were masked. We have combined holographic stimulation with MEA recordings (54), providing high spatial stimulation resolution of 8 µm spots. Prior to holographic stimulation, we took fluorescence images to visualize and manually map individual ChR2-EYFP-expressing neurons across the electrode grid. Individual neuronal positional information facilitated automatic guidance of the holographic stimulation from one targeted neuron to another. An episode of holographic stimulation of at least 20 pulses of 50 ms blue laser light (450 nm, 0.15 W/mm^2^) at 2 Hz (Figure 3A-B) was presented to each neuron. We were able to include on average 400 brightly ChR2-EYFP-labeled individual neurons per MEA (Figure S6). We detected robust light-induced activity by holographic stimulation (Figure 3).

**Figure 3.**
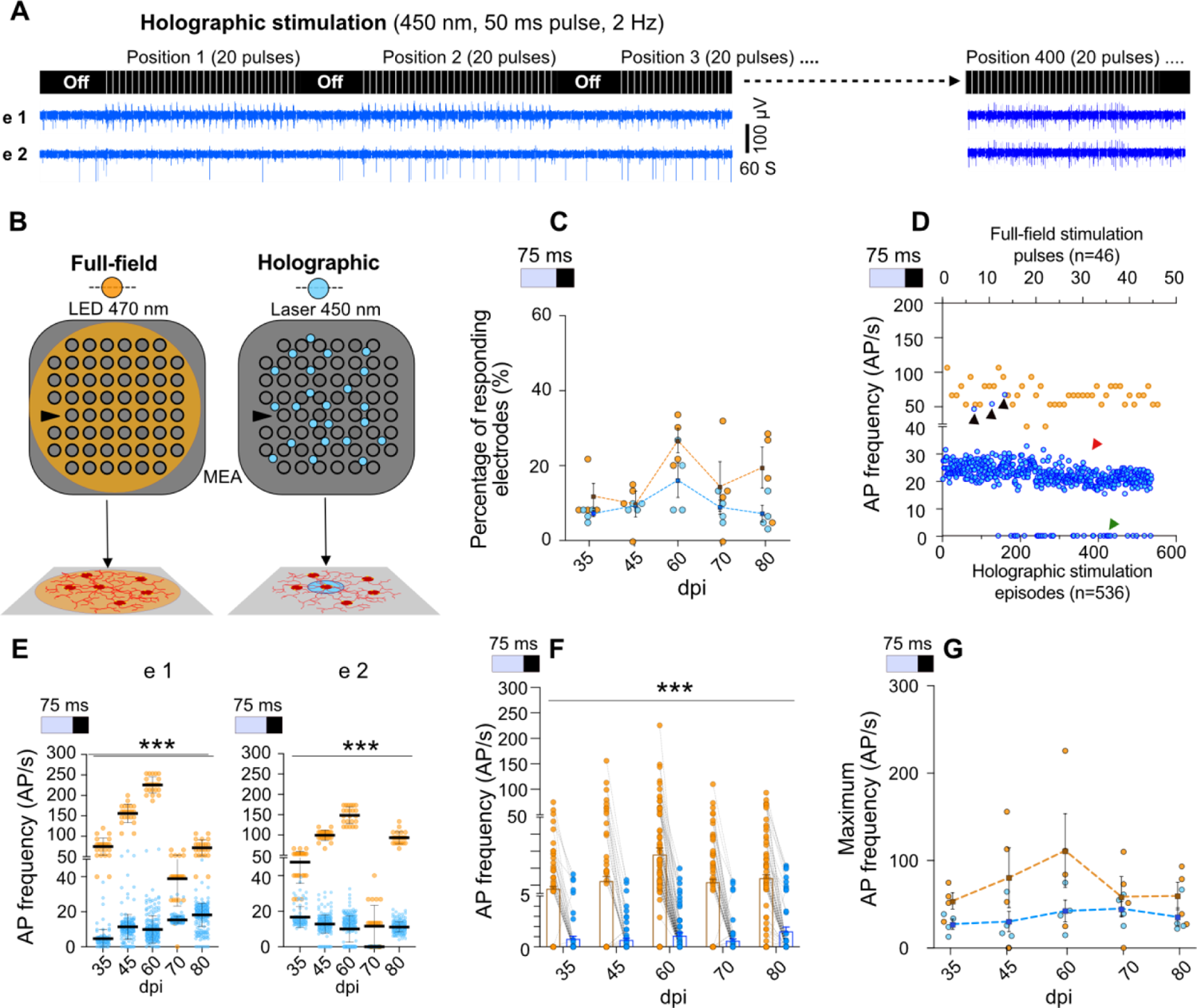
Activity evoked by holographic (blue) vs. full-field (orange) stimulation. (A) Holographic stimulation: neurons were detected by fluorescence microscopy and then stimulated 20-30 times during each holographic stimulation episode. Every holographic experiment at each dpi targeted around 400 neurons, each stimulated by an episode (upper panel). Sample profiles of activity in two electrodes (electrode 1 and electrode 2) show distinct responses to holographic stimulation of neurons at different positions in the network (lower panel). (B) Schematic of full-field stimulation affecting the whole network vs. holographic stimulation targeting individual neurons. (C) Percentage of electrodes responding to full-field or holographic stimulations in 4 different MEAs (each circle represents one MEA). (D) AP frequencies recorded during 536 subsequent holographic stimulation episodes (blue circles) or 46 pulses of full-field stimulation (orange). Black arrows indicate 3 holographic episodes that induced higher-frequency APs. Red arrow indicates moderate responses for the majority of episodes and green arrow shows episodes in which no response was recorded by an electrode. (E) Response in two single electrodes (left and right graphs) at different culture ages. (F) Average response to stimulation over time (N=4 MEAs, n=95 active electrodes). Average activity at each individual electrode in response to full-field and holographic stimulation are connected by gray lines. (G) Maximum firing rate in 4 MEAs. Mean ± SEM for each stimulation method at specific time points is illustrated as a thick horizontal line and error bars. In all cases, data from the two methods was compared using mixed-effect analysis followed by Sidak’s multiple comparisons test (***p<0.001).

Next, we compared the performance of full-field versus holographic optogenetic activation of iNGN neurons within the same MEA culture (N=4) over different time points of neuronal maturation. To exclude spontaneous baseline activity in between stimulation pulses, only the activity within the first 75 ms from stimulus onset (Figure 3) was considered in our analysis. There was no significant difference in the percentage of electrodes which responded to holographic vs. full-field stimulation (Figure 3C). However, the average AP frequencies triggered by holographic stimulation were significantly lower than with full-field stimulation (p<0.001; Figure 3E-F). A direct comparison of individual electrodes indicated that holographic stimulation led to lower AP firing rates of iNGN neurons (p<0.001; Figure 3E-F). The long-term trend of holographically-evoked AP frequencies differed between electrodes, as demonstrated by two sample electrodes in Figure 3E, one with stimulated AP frequencies increasing over time, the other with decreasing AP frequencies. Full-field stimulation resulted in increasing AP frequencies until day 60 in all electrodes, followed decreasing AP frequencies until day 80. (Figure 3E and Figure S7).

The average AP frequencies remained constant over consecutive episodes of holographic stimulation (Figure 3D and Figure S7). Within 400 holographic stimulation episodes, we detected two types of responses: moderate- and high-frequency APs. While high AP frequencies were rarely detected, moderate AP frequencies were the predominant response (Figure 3D and Figure S7). The maximum AP frequencies evoked by holographic stimulation were within the similar range to full-field stimulation (Figure 3D and Figure 3G).

Holographic stimulation did not evoke any network activity comparable to full-field stimulation. However, in holographic stimulation at the single-neuron level we were able to induce a similar intensity of responses to full-field stimulation. Additionally, holographic stimulation enabled the differences in per-electrode long-term AP frequency trends to be detected: these were obscured in data obtained by whole-network full-field stimulation.

### Distance-dependent responses of neurons to holographic stimulation

In random hiPSC-derived neuronal networks, only very few neurons are located close to an MEA recording electrode and are therefore accessible for direct recordings. Optical targeting of individual neurons by holographic stimulation, however, includes spatial information. It is possible to extract post-stimulus time histogram (PSTH) features with regard to the physical distance between stimulated neurons and recording electrodes from our data. We targeted around 400 neurons with holographic stimulation episodes over a period of 45 days (35-80 dpi). We analyzed data from six MEA electrodes that were consistently activated by holographic stimulation at different distances between the electrode and the stimulation site. For each episode, Euclidean distances between the targeted neuron and six active electrodes were calculated to correlate response features to physical distances over the entire MEA area (Figure 4A and Figure 4B). Distances were sorted into several intervals (Figure 4A and Figure 4B). For each holographic stimulation episode, the frequency of the PSTH peak at each electrode determined the evoked activity level. This data was pooled from all applied holographic episodes at different ages of the same network (n=1659 episodes). The latency of the PSTH primary peak from stimulus onset determined the time an evoked response of a neuron required to reach the active recording electrode. Holographic stimulation of nearby neurons, <30 µm, evoked significantly higher PSTH amplitudes (p<0.01 vs. larger distances, Figure 4C). These responses were associated with direct neuronal responses to the stimulation (Figure S4). We also observed a shorter PSTH peak delay when the stimulated neurons were less than 30 µm away from the recording electrode (p<0.01 vs. all other distances, Figure 4D). A shorter delay was also attributed to a direct neuronal activation, as shown by a correlation analysis of high PSTH peak amplitudes with short latencies (Figure 4E). Direct neuronal responses to holographic stimulation were mostly recorded within 30 µm of the electrode: lower PSTH amplitudes and larger latencies corresponded to distances longer than 30 µm and, therefore, reflected indirect responses through poly-synaptic connections (Figure 4C-E).

**Figure 4.**
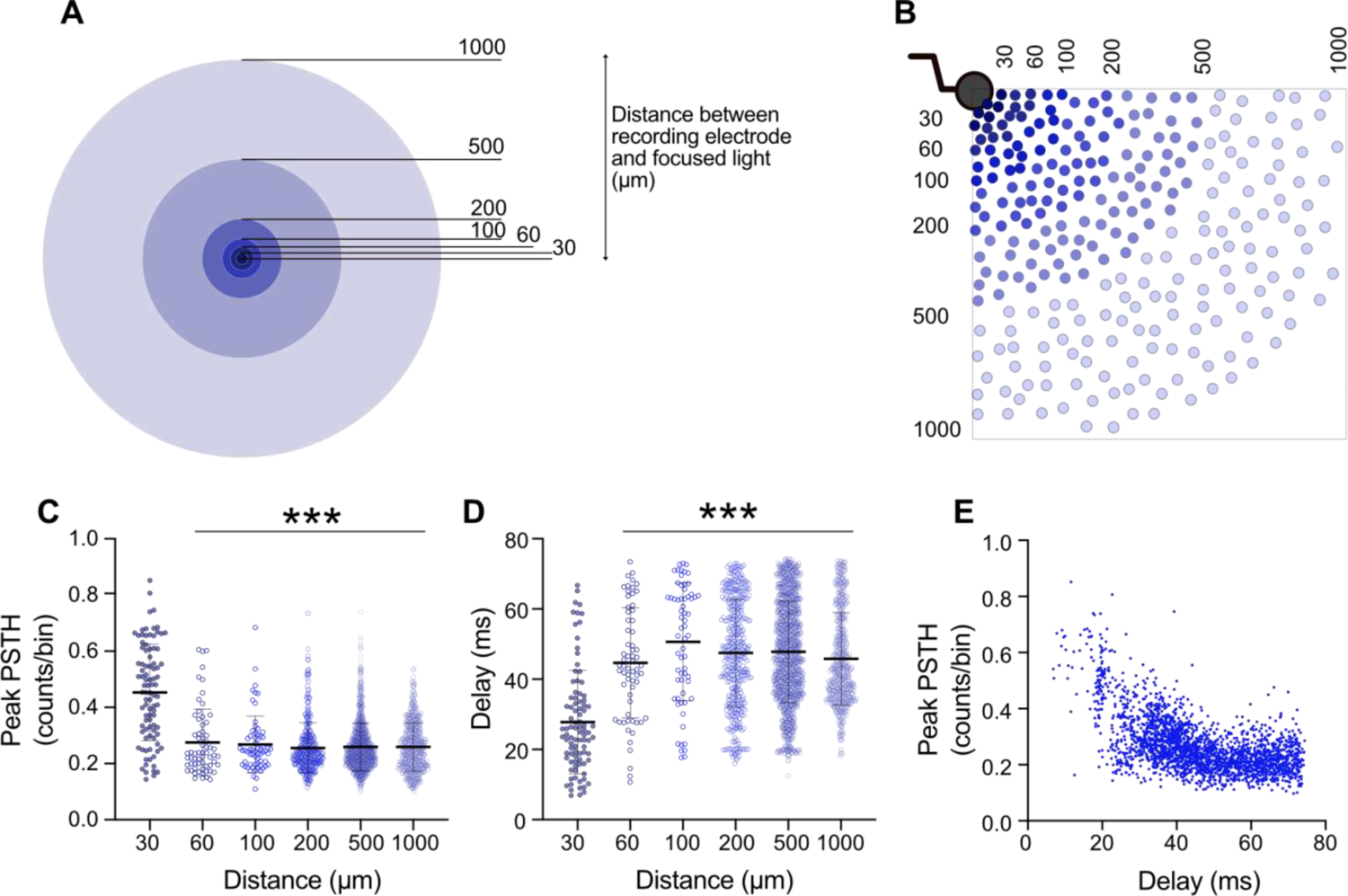
Distance-dependent direct and indirect responses to holographic stimulation. (A) Categorization of distances between each holographically stimulated neuron and each of the six analyzed electrodes was based on their radial distances from the edge of the electrode. (B) Sample schematic depicting distribution of stimulation points (ChR2-EYFP-expressing neurons) in relation to an electrode. (C-D) For each target neuron, 20 pulses were applied (450 nm, 50 ms, 2 Hz) and PSTHs were measured at six active electrodes. PSTH peak value and measured delay from stimulus onset were extracted and plotted based on categorized distances. Number of PSTHs studied at distances of 30, 60, 100, 200, 500, and 1000 µm were 105, 67, 65, 379, 1276, and 880, respectively. Each dot represents PSTH peak value (C) or peak position (D) for individual stimulation episodes. Average values represented as mean ± SEM with thick horizontal lines and vertical error bars. Mixed-effect analysis followed by Sidak’s multiple comparisons test was applied to compare between different distances. ***p<0.001 vs. the mean value at 30 µm distance. (E) Correspondence between individual PSTH peak values (amplitude) and PSTH peak positions (delay). Data were collected from recordings from one MEA, 35-80 dpi.

### The dynamics of connectivity motifs over time

Several methods have been proposed for extracting functional connectivity patterns from baseline MEA recordings (63, 64). For standard MEA recordings, however, many neurons in the network are beyond the typical recording distance of the electrodes (Figure S1). Their contribution to the network structure is therefore not directly observable. Directly stimulating individual neurons facilitated the extraction of whole-network functional connectivity maps between stimulated and recorded neurons as a superposition of all the connections found (Figure 5A and Figure 5D). We collected connectivity data from different electrodes (n=17) that showed a validated stimulus response to at least one of the approximately 400 target neurons per MEA (N=4) (Figure 5D). For comparison, we extracted electrode-electrode functional connectivity from baseline recordings using spectral entropy ((64), Figure 5B) and from single-neuron holographic stimulation recordings (Figure 5C). For this representation, the connection strength between two electrodes was measured as the sum of simultaneous responses to one stimulation episode, normalized to the overall number of stimulation episodes. This analysis was crucial to determine the differences and similarities between functional connectivity maps. Connectivity maps from cultures of different ages showed noticeable changes, including shifts in functionally connected electrodes and changes in the number of electrodes connected to each targeted neuron (Figure 5A and Figure S8).

**Figure 5.**
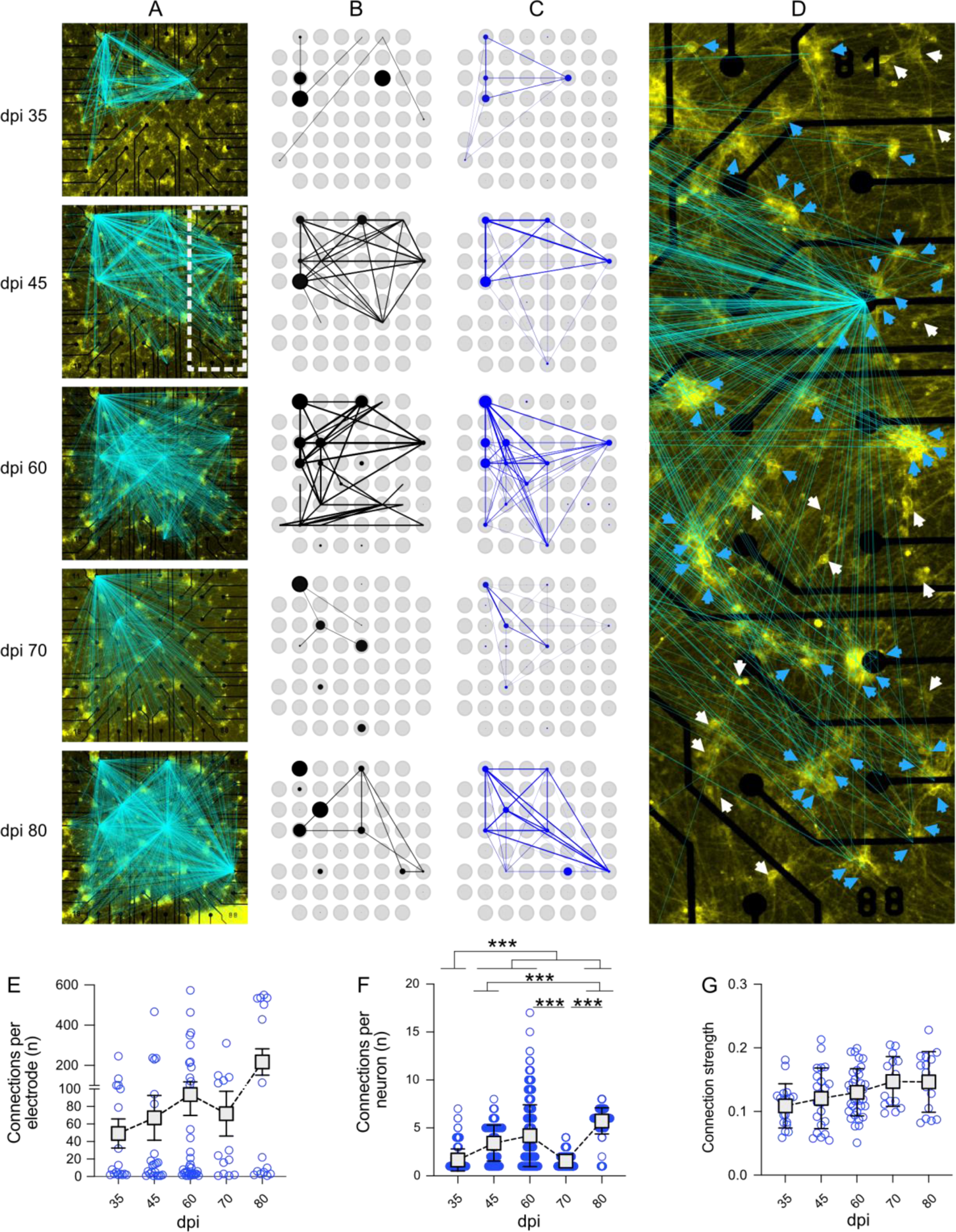
Functional connectivity maps over time based on spike-clustered data. (A) Development of the number of validated connections between stimulated neurons and recording electrodes over 80 dpi. Lines indicate a valid connection and are overlaid on the fluorescence image of the sample on the day of recording. Line intensity scales with connection strength. (B) Functional connectivity of the sample calculated for baseline activity and (C) holographic stimulation. Thickness and transparency scale with connection strength. Blue and black circles show general activity of the respective electrode, scaled to the maximum for each mode of experiment. (D) Shows a magnified view of the sample at 45 dpi (marked as white rectangle in (A)). White arrows indicate neurons which were stimulated but did not show a significant response, whereas blue arrows indicate neurons for which stimulation elicited a significant response. (E) The average number of neurons connected to an active electrode, normalized to the overall number of stimulated neurons per experiment (N=4 MEAs and n=56 active electrodes). (F) Number of neurons showing a valid response to the stimulation of a target neuron on each MEA (N=4 MEAs). (G) Strength of the connection of a neuron to the target neuron (N=4 MEAs and n=56 electrodes). Connection strength is measured as the maximum of the cross-correlation of the stimulus response of one neuron with the stimulus, represented by a train of boxcar functions. Data were compared between days using Kruskal-Wallis test followed by Dunn’s multiple comparisons test. *p<0.05, **p<0.01 and ***p<0.001 vs. specified date.

These connectivity maps of developing iNGN networks (N=4) at different ages were used to understand changes in connectivity parameters over time (Figure 5E-G). We counted the total number of neurons functionally connected to each active electrode (n=56) at different days. The average number of neurons connected to each active electrode (n=56 electrodes) increased from 49.18±16.64 at 35 dpi to 217±65.74 at 80 dpi (Figure 5E), although developmental age differences were not significant (p>0.99). The average number of functionally connected electrodes to each target neuron (N=4 MEAs and n=432 to n=868 target neurons depending on day of experiment) increased from 1.65±0.05 at 35 dpi to 4.19±0.11 at 60 dpi (p<0.001), followed by a decrease at 70 dpi (p<0.001 vs. 60 dpi), and reached its peak at 80 dpi (5.72±0.05, p<0.001 vs. 60 dpi, Figure 5F). The connection strength measured as the correlation coefficient between the stimulated spike train of spike-sorted APs and the stimulus signal slightly increased with culture age but this was not statistically significant (Figure 5G).

Connectivity parameters of developing networks extracted from holographic stimulation revealed that new aspects of iNGN network functional development, such as connection strength, do not necessarily follow increased or decreased network activity levels over time. Tracking multi-parametric network functional features with holographic stimulation extends our understanding of network functional development.

### Measuring connectivity motifs by precise holographic stimulation

Finally, we aimed to extract subnetworks within the random neuronal networks by tracking propagating evoked neuronal activity across the entire MEA area. Nearby and distant electrodes which indirectly received propagated neuronal activity, were extracted from MEA data by PSTH latency analysis at 34 dpi. When we simultaneously used full-field stimulation on all ChR2-expressing neurons, all latency-related information was lost (Figure 6B). In response to the full-field stimulation, different electrodes showed almost identical PSTH peak positions, as each electrode recorded direct neuronal activity at least from one ChR2-activated neuron (Figure 6B). However, by holographic stimulation of individual neurons, we detected that the PSTH profiles varied between neurons depending on direct or indirect activation by a holographic stimulus (Figure 6A and Figure 6C-D). To this end, we focused on three electrodes in which a holographic episode activated a neuron close to electrode 24 (Figure 6C) and evoked a two-component peri-event histogram profile. A direct response was measured during the light pulse (Figure 6C), followed by a delayed indirect response. This holographic episode induced a delayed response at electrode 83 (histogram peak between 50 ms and 500 ms) but did not affect the activity at electrode 45. Another holographic stimulation episode targeted a neuron close to electrode 45 and evoked a direct neuronal response at this electrode (Figure 6D). This episode also evoked an indirect response at electrodes 24 and 83 (Figure 6D). Based on this collective PSTH profile of electrodes 24, 45, and 83 in response to the holographic stimulation of two individual neurons close to electrodes 24 and 45, a functional connectivity motif within this random neuronal sub-network was extracted (Figure 6E).

**Figure 6.**
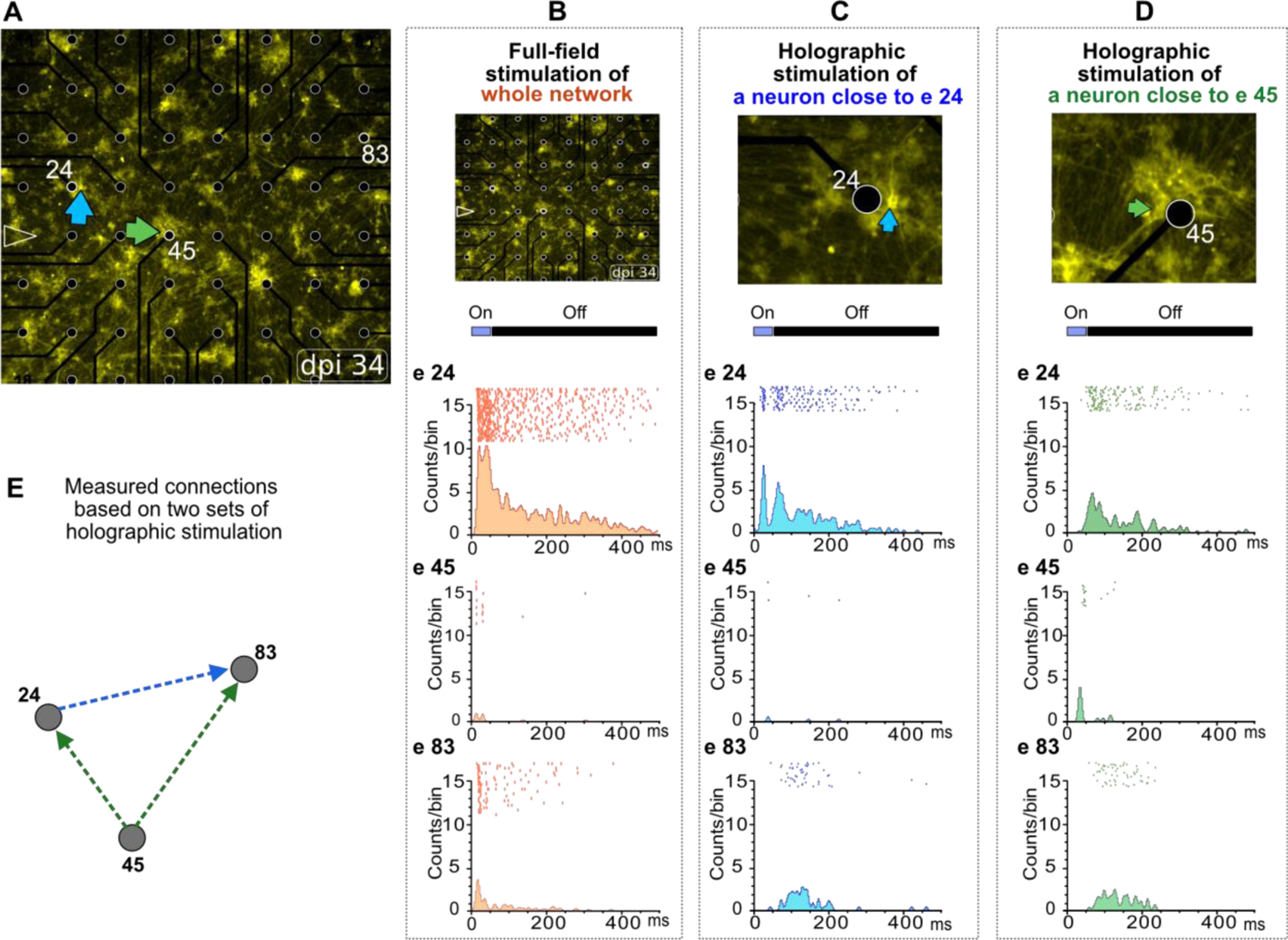
Representative connectivity pattern using PSTH profiles upon holographic stimulation. (A) Network morphology at 34 dpi with arrows indicating two neurons close to electrodes 24 and 45, which were stimulated separately by two episodes of holographic stimulation (n=20 pulses per episode) or simultaneously by full-field stimulation (n=44 pulses). (B-D) Responses to full-field stimulation (B) and holographic stimulation of a single neuron (C, D) in electrodes 24, 45, and 83 are shown. Upper panels show microscopy images of the whole network (B) or the stimulated neurons close to electrodes 24 (C) or 45 (D) (marked with blue and green arrow, respectively). Lower panels show peri-event rasters and histograms of the evoked activity in electrodes 24, 45, and 83 in response to full-field (B) or holographic (C, D) stimulation. (E) Schematic of connectivity pattern between three electrodes extracted from responses to holographic stimulation of two neurons shown in (C) and (D).

In conclusion, single-neuron holographic stimulation enabled propagating activity to be tracked across sub-networks of neurons based on PSTH timing in order to detect defined connectivity motifs.

## Discussion

To date, most studies on deciphering functional properties of hiPSC-derived neuronal networks have only captured early developmental time points. In addition, only basic functional features such as neuronal firing rates have been studied (25,65–67). Here, we combined time-lapse functional MEA studies of developing human stem-cell-derived neuronal networks with full-field and holographic optogenetic stimulation platforms, providing access to multiple functional features over time. Key to our study was the establishment of a robust long-term neuronal network culturing system, facilitating stable MEA recordings over months. The two-step protocol, first inducing neurogenesis on matrigel-coated surfaces, followed by re-seeding of neuronal precursors on poly-D-lysine (PDL)-coated MEAs, was crucial to provide long-term neuronal adhesion to the MEAs. We further supported the neuronal cultures with specialized commercial media, which was essential for neuronal maturation and function. In addition, we integrated an established astrocyte co-culture system in which the astrocytes were placed in close proximity to, but still separated from, the neurons (Figure 1). These glial cells secreted additional trophic factors into the media that supported long-term neuronal survival and maturation (28). The beneficial role of astrocytes on hiPSC-derived neurons is well established, and was demonstrated here by increases in firing rates, dendritic arborizations, and maturation of excitatory glutamatergic synapses (35). The astrocyte coverslip further shielded the neurons placed on the electrode array area from shear forces during media exchanges (22), and thereby provided a stabilized microenvironment. Media evaporation resulting in pH and osmolality changes represented another challenge for long-term electrophysiology recordings outside of the incubator (68). The lids prevented this, and facilitated long holographic stimulation sessions of the same MEA preparation over the time course of functional network development.

Our recorded spontaneous activity data are in line with previous studies using hiPSCs in which neurogenesis was triggered by Neurogenin-1 and −2 overexpression (25). Sparse APs were detected from 25 dpi onwards, and developed into burst activity, reaching a peak around 60 dpi followed by a slight decrease in activity at later time points. Developing primary cortical and hippocampal networks have shown comparable trends in activity over time *in vitro*, but these cells were already active at 14 days *in vitro* (div) and a peak activity was detected at 21 div, followed by decreased activity levels at later time points (23,61,69). Hence, the development of iNGNs, and also other human stem-cell-derived neurons (59), very likely reflect similar maturation patterns regarding neuronal activity. However, human neuronal cultures require longer time periods to become fully functional. Our previous work, based on whole cell patch-clamp recordings from iNGN neurons at different developmental time points, showed a trending decrease in resting membrane potential (RMP) up to 35 dpi (−75 mV) (28). The decrease in RMP was concomitant with increased expression of voltage-gated sodium and potassium channels and elevated spontaneous postsynaptic currents (sPSCs) (13, 28). While our previous patch-clamp study required different neuronal preparations for each recording day, our MEA study on identical cultures over time corroborated all findings on *in-vitro* human neuronal network maturation. Burst activity is mainly derived from contributions of excitatory and inhibitory synapses (62, 70). Spontaneous burst activity in iNGN networks appeared at around 35 dpi and reached its peak at 60 dpi. In hiPSC-derived brain organoids, highly synchronous activity has also been reported at around day 60: this later changes to rhythmic activity (71).

Instead of just waiting until spontaneous neuronal activity started to occur, we applied optogenetic stimulation. Indeed, we detected robust neuronal activity at earlier time points, which is in line with previous studies using electrical stimulation for hiPSC-derived (18, 72) and optogenetic activation of mouse ESC-derived neurons (73). We further showed that full-field stimulation evoked a combination of direct and delayed PSTH peaks, highlighting poly-synaptic connections that were blocked by AMPA- and NMDA-receptor antagonists (Figure S4). Optogenetic stimulation served as a booster for hiPSC-derived network maturation (74, 75). Still, further functional studies are required to explore the optimal continuous optogenetic stimulation parameters to accelerate functional maturation of human stem-cell-derived neurons. In our study, the neuronal cultures were stimulated by light during the recording sessions. Testing continuous and regular light stimulation within the incubators might further accelerate neuronal network maturation. This intervention will significantly decrease the human neuronal culturing period, saving valuable resources.

Our holographic stimulation platform revealed functional subunits within neuronal networks better than full-field stimulation. When all ChR2-expressing neurons were activated simultaneously by full-field stimulation, connectivity patterns between neurons were mostly masked. Holographic stimulation of ChR2-expressing neurons enabled single-cell activation, which allowed us to extract the propagating neuronal activity within sub-networks. For semi-automated holographic stimulation, on average 400 neurons were marked manually, followed by automated stimulation using a programmable pattern. To decrease the duration of the holographic stimulation sessions we increased the frequency of the applied stimuli to 2 Hz compared to 0.5 Hz full-field stimulation pulses. Responses to the applied stimulus frequency were robust and reproducible (Figure 5). Each episode of holographic stimulation targeted one selected neuron. The corresponding evoked activity was evaluated using the Kolmogorov-Smirnov test and used to identify connections to target neurons across the electrode area. These connections were used to extract overall network connectivity maps and connection strengths by cross correlations to the stimulus signal. We detected dynamic connectivity patterns: the total number and strength of functional connections increased with increasing culture age, and was dependent on the network activity levels. As a proof of principle, repeated stimulation sessions at different culture ages provided insights into the network connectivity dynamics of developing networks. It was possible to extract connectivity motifs by evaluating PSTH peaks and the latency of single-neuron stimulation.

So far, connectivity studies have only been performed on developing primary cortical and hippocampal circuits using spontaneous activity data (76–78) and with hiPSC-derived networks based on electrode-to-electrode functional connections (59). However, the latter method excluded a major part of the network activity, as most neurons were located between electrodes and therefore not accessible (79). Holographic stimulation made correlating electrode activity to neurons located far from electrodes possible, and thus increased the spatial resolution of the connectivity maps.

## Conclusion

Overall, we demonstrate that holographic optogenetic stimulation of human neurons is robust and superior to full-field stimulation for extracting additional detailed neuronal properties. In the present work we applied MEA electrophysiology based on standard transparent MEA devices that offer 8×8 electrodes (79). Next, we will also integrate MEAs with higher electrode density, such as CMOS-based MEAs, into our holographic stimulation platform. In this way, read-out efficiency will be improved and further details of the network function will be revealed (77). Our holographic stimulation platform is also capable of targeting several neurons simultaneously by generating multiple foci (54). In addition, the flexibility of the system will be further improved by integrating combinations of the optogenetic actuators, and controlling different neurons with diverse wavelengths.

These technological advances have extended the competence of the holographic stimulation for deciphering the functional features of developing hiPSC-derived networks: this will serve both basic biomedical research as well as studying pathophysiological features of developing networks derived from human patients. Having fully-functional human neurons is an important step to guide this model system out of its infancy, and enable it to become accepted as a complementary platform to animal research within the neuroscience community.

## Materials and Methods

### Long-term hiPSC-derived neuronal culture on MEA chips

We adapted a two-step cell-seeding protocol that included short-term induction of iNGN cells for 5 days in a Matrigel (Corning)-coated well plate followed by re-seeding differentiated neurons on PDL (Merck)-laminin (Sigma)-coated MEAs (Multi Channel Systems). Human-derived iNGN neurons (Busskamp et al., 2014) were thawed and passaged twice before seeding them in Matrigel-coated well plates. These cells were cultured in mTeSR™1 medium (mTeSR™1 Basal Medium with mTeSR™1 5× Supplement (Stemcell Technologies) and 1% penicillin-streptomycin (P/S; Thermo Fisher Scientific)). For neural induction, iNGN cells were treated with 0.5 μg/ml doxycycline (Sigma) for 5 consecutive days (Figure 1). To initiate the expression of EYFP-tagged ChR2 in iNGN cells, a lentiviral vector (plv-ef1a-ChR2-EYFP) was added one day after seeding within a biosafety level-2 facility. Undifferentiated cells were removed by adding Ara-C (5 μM final concentration; cytosine β-D-arabinofuranoside hydrochloride, Sigma) at day 4 post-induction (4 dpi). Standard MEA chips were plasma treated for 2 minutes (ambient air, 0.3 mbar), coated with PDL (50 µl per electrode area in 0.1 mg/ml final concentration) and incubated overnight at 37°C. MEAs were washed with sterile deionized water (diH_2_O) (3 times, each time 10 min) and dried under the hood with laminar flow. Laminin (50 µl; 0.05 mg/ml concentration) was added to the PDL-coated MEAs and incubated for 3 hours before reseeding the 5 dpi iNGN cells, which were dissociated by incubating them with Accutase (Sigma) for 5 minutes. The dissociated cell suspension was centrifuged (1400 rpm, 4 min), supernatant was removed and re-suspended in complete BrainPhys™ (BrainPhys™ Neuronal Medium (Stemcell Technologies)+ 1% P/S + NeuroCult™ SM1 Neuronal Supplement (Stemcell Technologies) + N2 Supplement-A + 20 ng/ml recombinant human BDNF (Peprotech) + 20 ng/ml recombinant human GDNF (Peprotech) and 200 nM ascorbic acid (Sigma)). Differentiated cells were re-seeded on PDL-laminin-coated MEA chips (around 100,000 cells per cm^2^ including electrode area).

Rat primary astrocyte cultures (Thermo Fisher Scientific, A1261301) were prepared in parallel (Figure 1) on PDL- and laminin-coated coverslips (0.1 mg/ml and 0.05 mg/ml, respectively). Paraffin dots were already placed on the coated side of the coverslips (Figure 1). Astrocytes were cultured in astrocyte medium including DMEM + 4.5 g/l d-glucose + pyruvate plus N2 Supplement, 10% One Shot™ fetal bovine serum and 1% P/S (all provided by Thermo Fisher Scientific). Astrocyte media was changed to complete BrainPhys media one day before re-seeding the neurons on MEA chips. Astrocyte cultures on paraffin coverslips were washed with PBS without calcium and magnesium (Thermo Fisher Scientific), flipped and placed on top of the re-seeded neural cells. Then, complete BrainPhys™ (BrainPhys™ Neuronal Medium (Stemcell Technologies) + 1% P/S + NeuroCult™ SM1 Neuronal Supplement (Stemcell Technologies) + N2 Supplement-A + 20 ng/ml recombinant human BDNF (Peprotech) + 20 ng/ml recombinant human GDNF (Peprotech) + 200 nM ascorbic acid (Sigma)) was added to the MEA rings, covering neuronal and astrocyte cultures (Figure 1). Half of the media was exchanged with fresh complete BrainPhys™ every week.

### Microscopy

An inverted EVOS FL Imaging System (Thermo Fisher Scientific) was used to prepare brightfield (10× and 20×) and fluorescent images (20×) from neuronal cultures on MEA chips at different days post-induction (Figure 1). High-resolution fluorescent images were used to detect neural cells expressing the EYFP-tagged ChR2. In all full-field and holographic light-stimulation experiments, high-resolution images were prepared from all cultures just before MEA recording. From each MEA culture, 20 images (20× magnification) were acquired to cover the entire electrode area (1.5 mm^2^). These images were then automatically stitched using ImageJ software (80), adjusted for their contrast, and used to target individual neurons in holographic light stimulation (Figure S1 and Figure S6).

### MEA electrophysiology

Neural network spontaneous and light-evoked activities were recorded using standard MEA chips (60 MEA200/30iR-Ti-gr) on an MEA1060-Inv-BC amplifier (sampling rate 25 kHz) using the MC_Rack software, all provided by Multi Channel Systems (MCS). In each MEA chip, 59 electrodes (8 × 8 matrix of electrodes including a counter electrode) simultaneously recorded extracellular activities from different parts of the network. Data acquisition was performed at a sampling frequency of 25 kHz. During the recording, temperature was controlled by a built-in temperature controller (TC02, MCS). The MEA chip and amplifier were mounted on an inverted microscope (Nikon Eclipse Ti). All recordings were performed in a dark Faraday cage shielding external electromagnetic and optical noises. Spontaneous activity was recorded from 15 dpi onward (Figure 1). Baseline spontaneous activity was recorded for 3 minutes before starting the full-field stimulation experiment on all recording days. Recording of spontaneous and evoked activities was performed in the presence of astrocyte culture.

### Full-field optogenetic stimulation

Before performing functional recordings, the center of each MEA chip (electrode area) was aligned with the center of the 640 nm light beam and its focal plane. Full-field stimulation and functional recordings were performed from 15 dpi (Figure 1). A spectra 4 (Lumencor) LED light source was used to apply full-field stimulation (470 nm, 50 ms pulses, 0.5 Hz) at 0.46 mW/mm^2^ intensity through a 4× objective lens of the inverted Nikon microscope. Blue light pulses were applied without including the filter cubes in the light path. The intensity of the light pulses was adjusted using commercially-available software (Lumencor Graphical User Interface). Stimulation protocols including pulse width, intervals (frequency), and number were controlled by a custom Python script and pCLAMP 11 software (Molecular Devices). Full-field stimulation light pulse properties including light-ON and light-OFF timestamps were digitized and recorded parallel to electrophysiology signals in MC-Rack recording software (MCS).

### Holographic stimulation

#### Holographic stimulation setup

The setup used has been reported previously and can be seen in Figure S9 (54). Collimated light from a laser diode (450 nm, Thorlabs) illuminates a ferroelectric liquid crystal on a silicon spatial light modulator (SLM) (QXGA-R9, 1536×2048 pixels, 8.2 µm pixel pitch, Forth Dimension Displays), which offers binary phase modulation at a maximum refresh rate of 400 Hz. Light reflected off the SLM is converted to true binary phase modulation using a linear polarizer. Fresnel Holograms displayed by the SLM are demagnified by a Keplerian telescope (magnification β = 25 mm/180 mm = 1/7.2). The minimum focusing distance for our setup according to (54) is 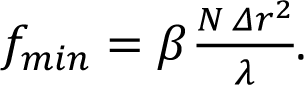 With N=1536 being the number of pixels along the shorter side of the SLM, Δr=8.2 µm being the pixel pitch of the modulator, and Δ=450 nm the wavelength of the laser, f_min_ ≈ 6.4 mm for this setup. Taking into account the focal length f_3_ of the lens facing the sample, the overall working distance is then d_w_ = f_min_ + f_3_ = 31.4 mm, which is required to fit the MEA recording device underneath the front optical element. The minimum focus diameter achieved was 8.8 µm. The system is calibrated and operated using a self-programmed graphical user interface (GUI) written in MATLAB. Co-planarity of the sample and the focal plane of the stimulation system was ensured by matching maximum contrast of MEA electrodes with a minimal focus diameter as observed by the microscope camera. Calibration of the setup is performed by projecting a user-defined set of foci onto the sample. The centers of these foci are then detected using a Gaussian fit function. Sample-plane coordinates of the foci are matched to the SLM-plane coordinates of the Fresnel zone plate centers, using a 2nd order polynomial fit to enable precise stimulation of individual neurons. To match coordinates of the high-resolution stitched fluorescence images to the lower-resolution camera images of the MEA system, the electrode grid is used to correlate both images. Centers of mass the electrode heads of both images are matched using a bilinear image transformation which accounts for rotation, shift, and linear scaling. This dual calibration makes it possible to calculate holograms which stimulate neurons in MEA camera coordinates which were previously selected in high-resolution fluorescence images.

#### Hologram calculation

Fresnel holograms were used for holographic stimulation. They were calculated by convoluting the desired illumination pattern on the sample with a pre-calculated Fresnel zone plate for the minimum focus distance f_min_, followed by either phase thresholding or bidirectional error diffusion of the phase for binarization (81), depending on the complexity of the hologram.

#### Holographic stimulation protocol

Before the experiments, individual neurons were selected for holographic stimulation from fluorescence images using our self-programmed MATLAB GUI (Figure 2). The GUI was also used to define the stimulation parameters including light pulse duration, frequency, and repetitions. After marking and saving the parameters, holographic stimulation was automatically guided from one targeted neuron to another until all marked neurons were stimulated. Each neuron received 20-30 pulses of 50 ms blue light (450 nm) at 2 Hz, which we call an episode of holographic stimulation (Figure 2). Depending on the network on each MEA and the number of available neurons, on average we applied around 400 episodes, each including one neuron at a particular time point (dpi).

### Inhibition of excitatory and inhibitory synapses

Two MEA cultures were used to block the excitatory glutamatergic synapses in the network at 110 dpi. We applied a combination of 2,3-dihydroxy-6-nitro-7-sulfamoyl-benzo[f]quinoxaline (NBQX; an AMPA receptor antagonist) and (2R)-amino-5-phosphonopentanoate (APV; a selective NMDA receptor antagonist). Before NBQX+APV treatment, spontaneous activity and response to full-field stimulation was recorded. Half of the medium in the MEA ring was drained and kept in an incubator. Then NBQX (10 µM final concentration) and APV (50 µM final concentration) were prepared in 500 μl of fresh BrainPhys™ medium and added to the rest of the medium in the MEA ring. After 5 minutes incubation, spontaneous activity was recorded in the presence of the NBQX+APV, and then full-field stimulation was applied at different light intensities (Figure S4). Later, NBQX+APV was washed out 2 times with warm complete BrainPhys™ medium. Then the pre-used media plus fresh media were mixed (1:1) and added to the MEA chamber. After 30 minutes incubation, activity was recorded again under non-stimulated and full-field stimulated conditions. Two other MEA cultures were selected to block inhibitory GABA A-receptors using gabazine (10 µM final concentration) at 115 dpi. The protocol for gabazine treatment was the same as for NBQX+APV. Spontaneous activity was recorded before adding gabazine and then in the presence of gabazine.

### Data analysis

#### AP frequency and burst frequency at baseline (spontaneous activity)

Long-term activity data was collected from 4 MEA chips with a total number of 95 active electrodes on seven different days (between 15 and 80 dpi). Recorded electrophysiology data was processed offline. Raw signals were filtered (Butterworth 2nd order, high-pass filter cut-off at 100 Hz) to remove low-frequency noise. The timestamps of the action potentials (APs) were detected by negative and positive thresholds (±5σ of the peak-to-peak noise). These timestamps were imported into Neuroexplorer software (Plexon Inc) to extract action potential frequency and burst features. For long-term spontaneous activity we included only active electrodes in the analysis. An electrode is considered active if it showed activity on at least two sessions (days) of recording and the AP frequency was more than 0.2 Hz in both sessions. Most active electrodes showed consistent activity during the whole experimental period. For the electrodes that showed activity only on some days the activity on all other days was set to zero: it was then possible to track the development of activity features over time.

Bursts were detected and extracted using Neuroexplorer software based on the following parameters: 20 ms maximum inter-spike interval to start the burst, 10 ms maximum inter-spike interval to end the burst, 10 ms minimum inter-burst interval, 20 ms minimum burst duration with at least 4 spikes in each burst (82). Different features of burst activity, including burst frequency, burst duration, and percentage of APs forming burst activity, were extracted.

#### AP frequency and burst frequency evoked by full-field stimulation

We applied an identical method of data extraction for collecting the full-field evoked responses from 95 active electrodes of 4 MEAs. We used the evoked activity between the first and last applied pulse of the full-field stimuli, including both the 50 ms pulse duration and 1950 ms pulse intervals (light-OFF periods between pulses; Figure 2). The APs and burst frequencies collected on each recording day were compared with baseline spontaneous activity using Wilcoxon matched-pairs signed rank test.

Baseline AP frequency and burst frequency between NBQX+APV-treated and untreated conditions were also compared using Wilcoxon matched-pairs signed rank test (two MEAs and 39 active electrodes). Light-evoked AP and burst frequencies were compared between NBQX+APV-treated and untreated conditions using the non-parametric Friedman test followed by Dunn’s multiple comparisons test (Figure S4).

#### Comparing activity profiles of full-field vs. holographic stimulation

To compare the evoked activity profile resulting from holographic and full-field stimulation, only APs that appeared during the 50 ms light pulses and 25 ms after (75 ms in total) were used. This range included all direct neuronal responses to the light stimulus (Figure 3). Each response to each full-field stimulation pulse represented one data point. For holographic stimulation, average response to 20-30 pulses of each episode represented one data point. Because we applied around 400 holographic stimulation episodes (Figure 3 and Figure S7) each day, around 400 data points were collected for each electrode (Figure 3). The percentage of responding electrodes, and the average and maximum frequency of evoked APs were compared between full-field and holographic stimulation data based on mixed-effects analysis of variance followed by Sidak’s multiple comparisons test (Figure 3).

#### Peri-event raster and time histogram

Peri-event raster and time histograms were created by counting the number of action potentials after the light stimulus (0-500 ms) and sorting them in 3 ms time bins. Sequences of the responses for repetitive light pulses of each holographic stimulation episode or 23 pulses of full-field stimulation were aligned and superimposed in time.

#### Post-stimulus time histogram (PSTH) for holographic stimulation

For the single-cell stimulation data presented in Figure 4 and Figure 5 we calculated PSTHs using self-programmed MATLAB scripts. All spikes occurring in a 75 ms window after the stimulation onset were extracted and sorted into 40 µs bins. The resulting distribution was smoothed by a convolution with a Gaussian function with a width of 3 ms as detailed in the equation below.

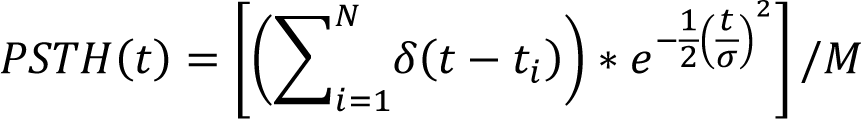

with *N* being the total number of spikes of one stimulation pulse occurring in a 75 ms window after stimulus onset, *t_i_* the arrival times of the individual spikes, 2σ = 3 ms the full 1/e width of the Gaussian function, *M* the number of stimulus repetitions per episode and ∗ the convolution operator.

PSTHs of valid connections (see Methods: Single neuron stimulation functional connectivity maps) were only considered if they consisted of more than 20 individual spikes. PSTH maxima were normalized to the number M of stimulations per episode.

Distances between stimulation site and recording electrode were calculated as the Euclidean distance between predefined stimulus location from the fluorescence microscope images and the centers of the recording electrodes.

#### Baseline functional connectivity maps

Baseline connectivity maps were extracted using the spectral entropy synchronicity measure (Kapucu et al., 2016). For further details on spectral entropy evaluation see supplementary material (Part S11). To avoid evaluating inactive electrodes, only signals with at least one event exceeding the threshold of 6σ peak-to-peak noise were chosen for further processing.

#### Single neuron stimulation functional connectivity maps

For the evaluation of connectivity using holographic single-cell stimulation, all episodes were evaluated separately. Detected spikes were sorted into separate units using wave_clus3 (83). To achieve consistent cluster detection, baseline and stimulation experiments were evaluated together and split again in post processing. To detect connections from stimulated to recorded neurons, deviations between stimulated and unstimulated states of single units were distinguished using the displaced impulses function (DIF) (84). Further details on DIF evaluation can be found in the supplementary material. DIF functions were tested for deviations from baseline using a Kolmogorov-Smirnov (KS) test. Values of p<0.01 were considered to be significantly influenced by stimulation, and therefore a valid connection to the recording electrode. To avoid the possibility of misinterpreting the results based on one burst in the time series, displaced impulses functions were calculated and tested with the KS test for all N_stim_ stimuli of one episode, excluding spikes belonging to a specific stimulus event, resulting in *N_stim_ individual tests for one episode*.

DIFs from electrodes without baseline activity (spikes only evoked by light stimulation) were tested against distribution functions with all values equal to zero. Only episodes passing these N_stim_ tests were finally considered to be a valid connection between the light-stimulated neuron and one unit at a recording electrode.

Connection strengths were calculated as follows: For a stimulus pulse train

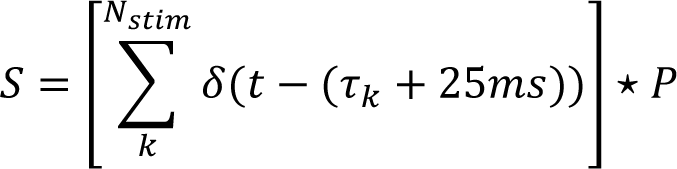

with the single pulse signal *P*,

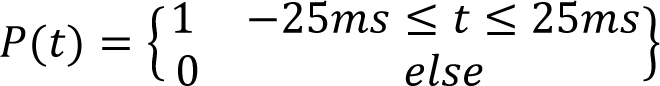

where *τ_k_* are the times of the stimulus onsets, and the connections strength is regarded as the maximum of the cross correlation of *S* and the spike events *E*

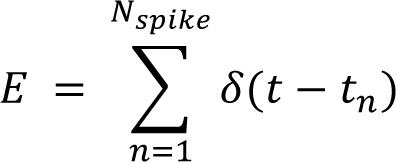

with *N_spike_* being the number of spikes in the time interval between the beginning of the first stimulus and the end of the last stimulus of one episode. For better overlap of *S, E* is additionally convoluted with a Gaussian function with a full 1/e width of 3 ms. However, because the overlap of the two functions is still very small, the connection strengths calculated this way are probably still very much underestimated.

So far, only connections between stimulated neurons and spike units on electrodes have been considered. To estimate functional connectivity maps in a similar way to the baseline connectivity maps, these connections have to be converted to electrode-electrode connections. To this end, all valid connections produced by one stimulation episode are considered. If there are valid connections from the stimulation site to a spike unit on more than one electrode, the connection strength between these electrodes is increased by one. Finally, connection strengths are normalized by the duration of the holographic stimulation experiment under the assumption that longer experiments with more stimulated neurons will, on average, have higher resulting connection strength values in a well-connected network. For comparison between days, all connection strengths are also normalized to the maximum value across days.

One main result of the holographic single-neuron stimulation is the set of all valid connections *n_ijk_k* of the neuron stimulated in episode i to the unit *j_k_* on recording electrode *k*. The number of functional connections per neuron is then:

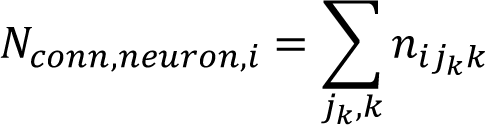

The number of valid connections per electrode, on the other hand, is then:

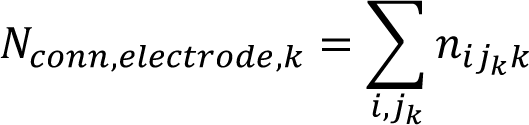

## Author contribution

Conceptualization: VB, JC, LB; Methodology: VB, JC, LB, FS, RH; Software: FS, RH; Validation: FS, RH, JS; Formal analysis: RH, FS; Investigation: RH, FS; Resources: VB, JC, LB; Data curation: RH, FS; Writing-Original draft: RH, FS, VB; Writing-Review & Editing; RH, FS, JS, LB, VB, JC; Visualization: RH, FS; Supervision; VB, JC, LB; Project administration: LB, VB, JC; Funding acquisition: JC, LB, VB.

## Acknowledgments

The authors thank Sara Oakeley for critical feedback on the manuscript. JC was supported by the Deutsche Forschungsgemeinschaft (DFG CZ 55/39-1 and the Reinhart Koselleck Project for High-Risk Research). VB acknowledges funding from the European Research Council (ERC-StG-678071– ProNeurons), the Deutsche Forschungsgemeinschaft (BU 2974/4-1, SPP2127 and EXC-2151-390873048–Cluster of Excellence– ImmunoSensation^2^ at the University of Bonn), the Pro Retina Foundation, the Paul Ehrlich Foundation, and the Volkswagen Foundation (Freigeist–A110720).

