## Supplemental Files for "Tracking long-term functional connectivity maps in human stem-cell-derived neuronal networks by holographic-optogenetic stimulation"

Title

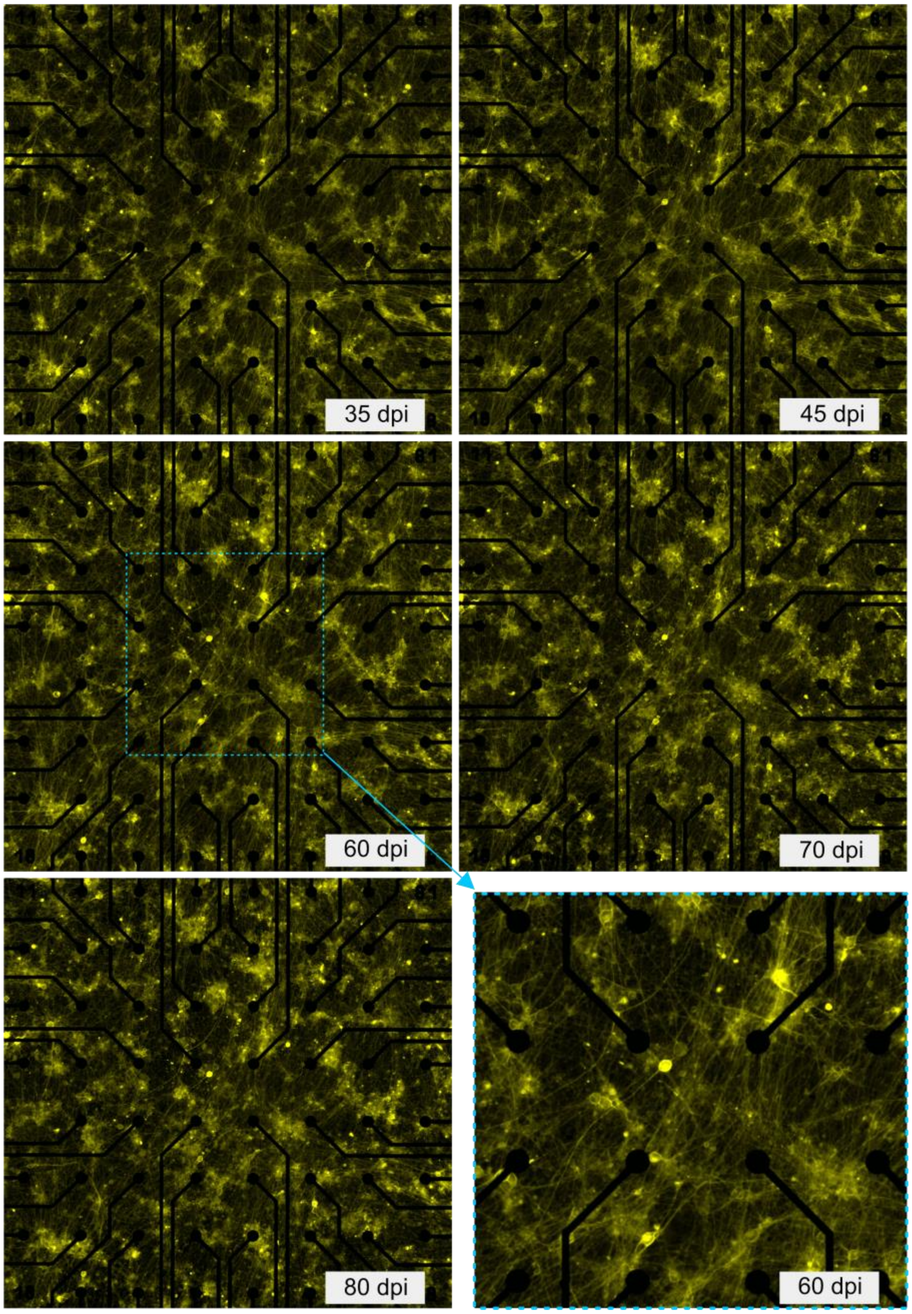


**Figure S1** Network morphology at different days post induction (dpi). Bottom right corner shows the magnified view of the network area covering 16 electrodes.

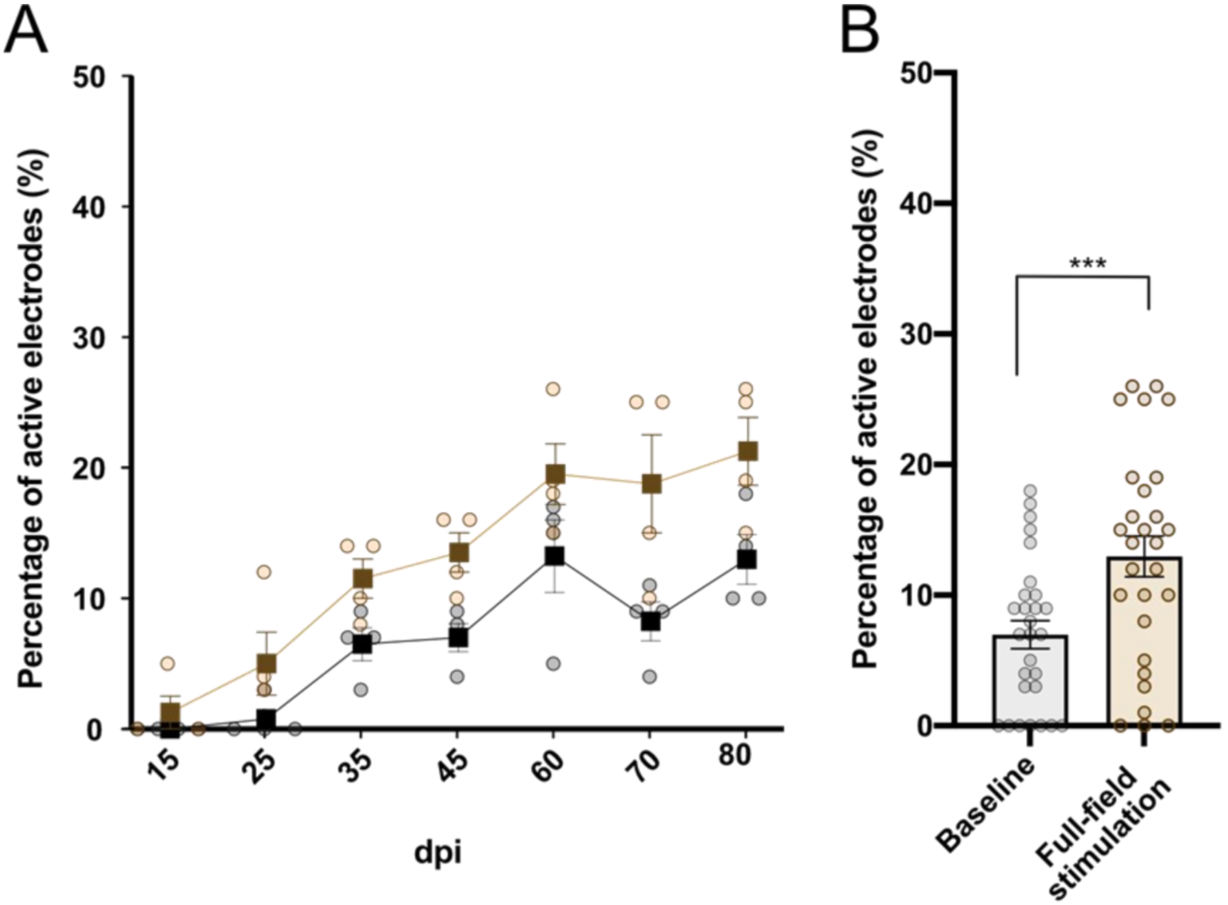


**Figure S2** Percentage of active electrodes at baseline and full-field stimulated conditions. (A) Percentage of active electrodes in 4 MEAs at different dpi. Mixed-effects model followed by Sidak's multiple comparisons test were applied to compare the groups in individual dpi. (B) Percentage of active electrodes pooled from 4 MEAs at different dpi. Wilcoxon signed-rank test was applied to compare percentage of active electrodes at baseline vs. full-field stimulation. ***p<0.001 vs. baseline.

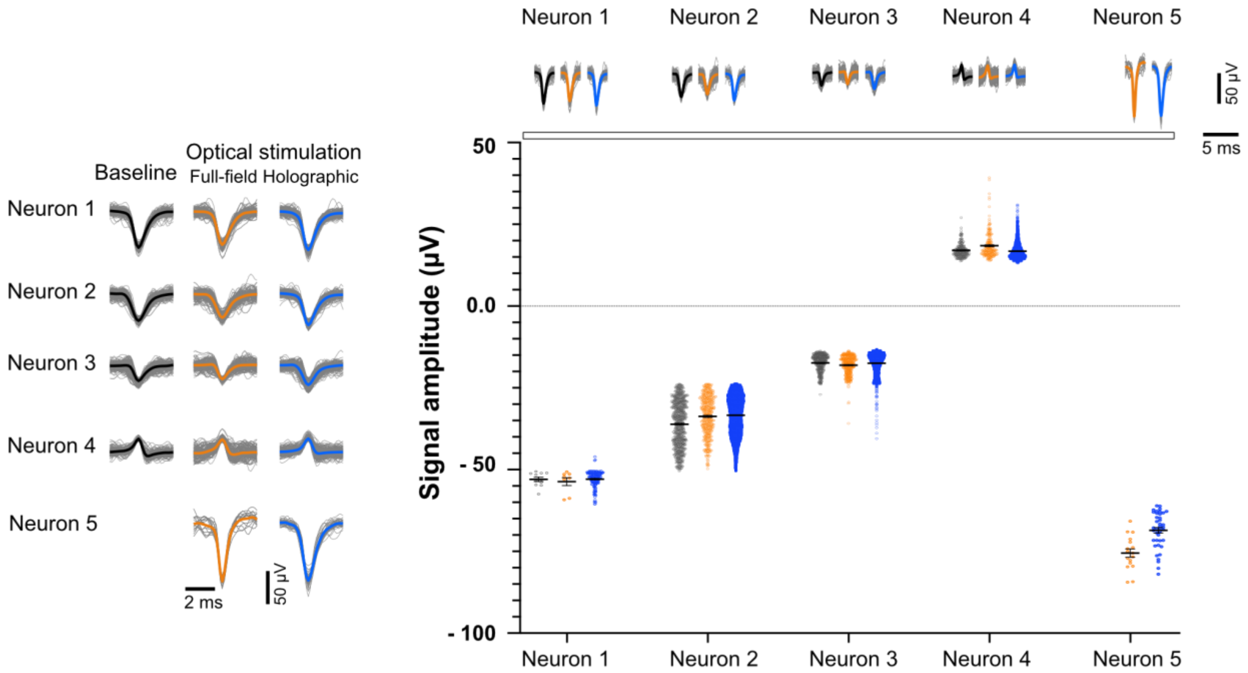


**Figure S3** Extracted number of active neurons from a single electrode at baseline, full-field, and holographic stimulation. (A) Each waveform represents activity recorded from one neuron. Neuron number 5 was only active during full-field and holographic stimulation. (B) Comparison of AP amplitude recorded from each neuron under the three conditions. There was no significant difference between waveform amplitude at full-field and holographic stimulation compared to the spontaneous activity. One-way ANOVA and Tukey's multiple comparisons test was applied to compare three pairs of waveforms at each neuron.

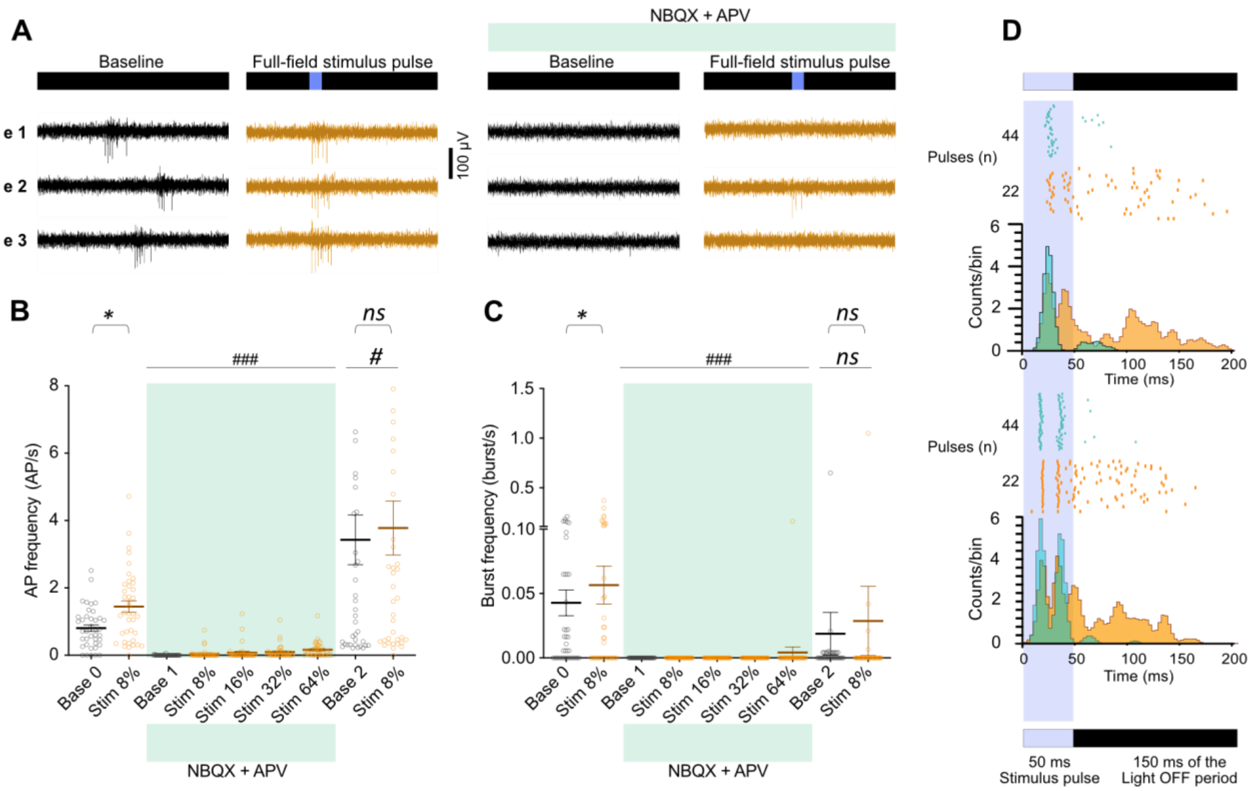


**Figure S4** Effect of AMPA and NMDA receptor antagonists on network activity features under baseline and optically-stimulated (full-field) conditions. (A) At 110 dpi, baseline spontaneous activity and evoked activity in full-field optical stimulation recorded before and after treatment with NBQX+APV and in the presence of NBQX+APV. Activity in three electrodes is shown at baseline (black) and under full-field stimulation (orange) before NBQX+APV treatment (left) and in the presence of NBQX+APV (right). Light stimuli (470 nm, 50 ms pulses at 0.5 Hz) with different intensities from 8% (0.46 $mW/mm^2$) to 64% (2.74 $mW/mm^2$) were applied. (B and C) Action potential frequency (left) and burst frequency (right) averaged across two MEAs only in active electrodes under different conditions. Each circle represents one electrode and each thick line represents the average activity under specific conditions. Baseline activity (Base) and responses to optical stimulation (Stim) before or after NBQX+APV treatment was analyzed using the non-parametric Wilcoxon matched-pairs signed rank test (*p<0.05 vs. baseline). The average response to the full-field stimulation in the presence of NBQX+APV is compared with Base 0 or Stim-8% before treatment using the non-parametric Friedman test, followed by Dunn's multiple comparisons test (#p<0.0, ###p<0.001 vs. Base or Stim-8% before treatment). (D) PSTH calculated 200 ms from stimulus timestamp in 3 ms bins in two electrodes (upper and lower graphs) in untreated (orange) and NBQX+APV treated (green) conditions. In each graph, the upper panel with raster plots shows the individual responses for 44 pulses or 22 pulses (arranged from bottom to top) in NBQX+APV treated (green) and untreated (orange) conditions, respectively. The resulting histogram has been plotted based on cumulative counts per bin.

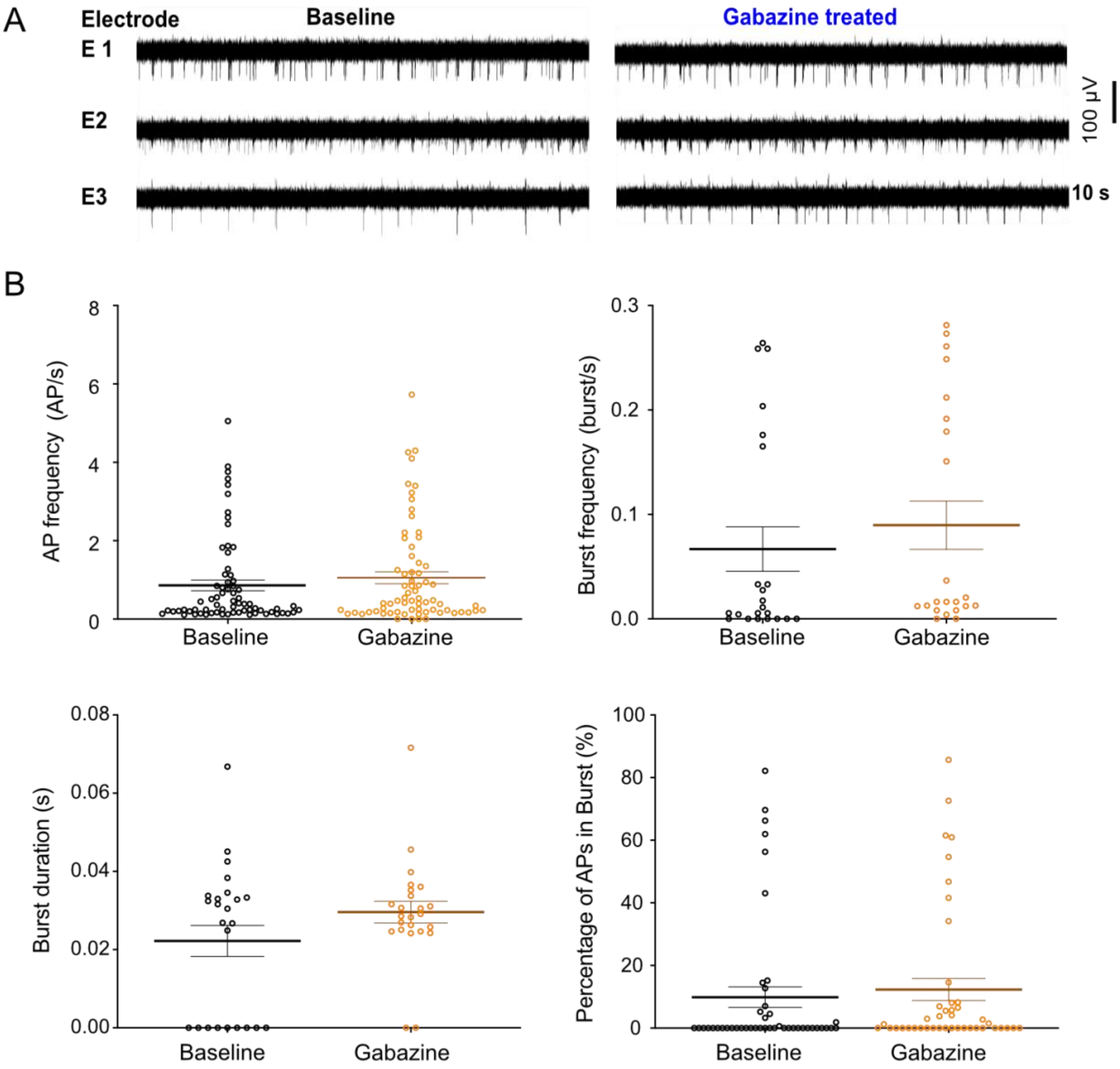


**Figure S5** Response of the iNGN-eDIO culture to inhibition of GABA-A receptors by gabazine. (A) Profile of the activity in 3 selected electrodes at baseline and in the presence of gabazine. (B) Action potential frequency and burst features were averaged across electrodes of 2 MEAs under treated and untreated conditions. Baseline activity and activity in the presence of gabazine was analyzed using the non-parametric Wilcoxon matched-pairs signed rank test (*$p<0.05$ vs. baseline). Data extracted from 2 MEAs at 115 dpi.

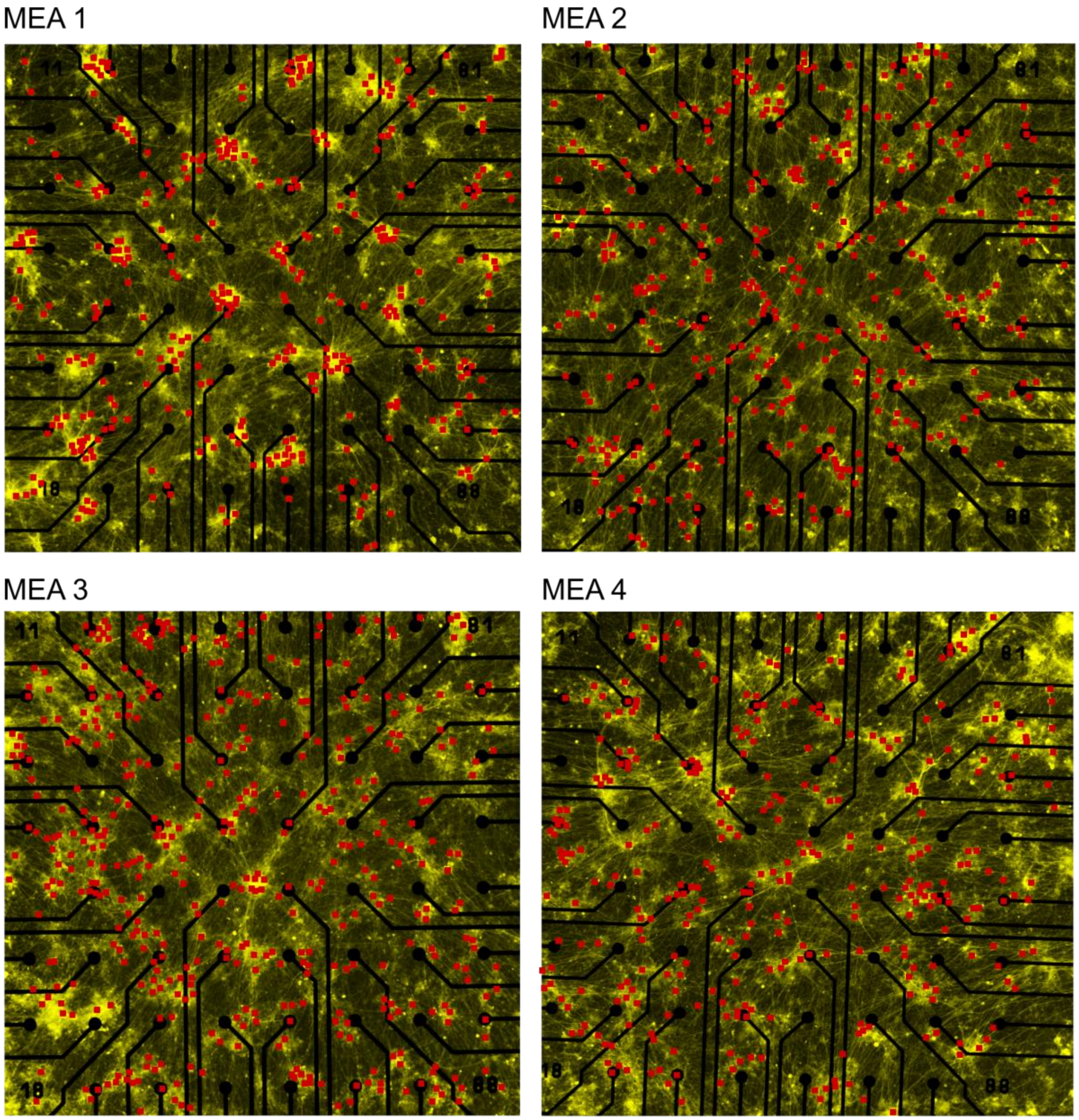


**Figure S6** Holographic stimulation points in 4 different MEAs at 60 dpi. Each red dot indicates a detected ChR2-EYFP-expressing neuron that has been stimulated by one episode of holographic stimulation (20 pulses, 450 nm, 50 ms, 2 Hz). Total number of applied holographic stimulation episodes varied between MEAs depending on the total number of neurons detected. Electrode pitch was 200 µm.

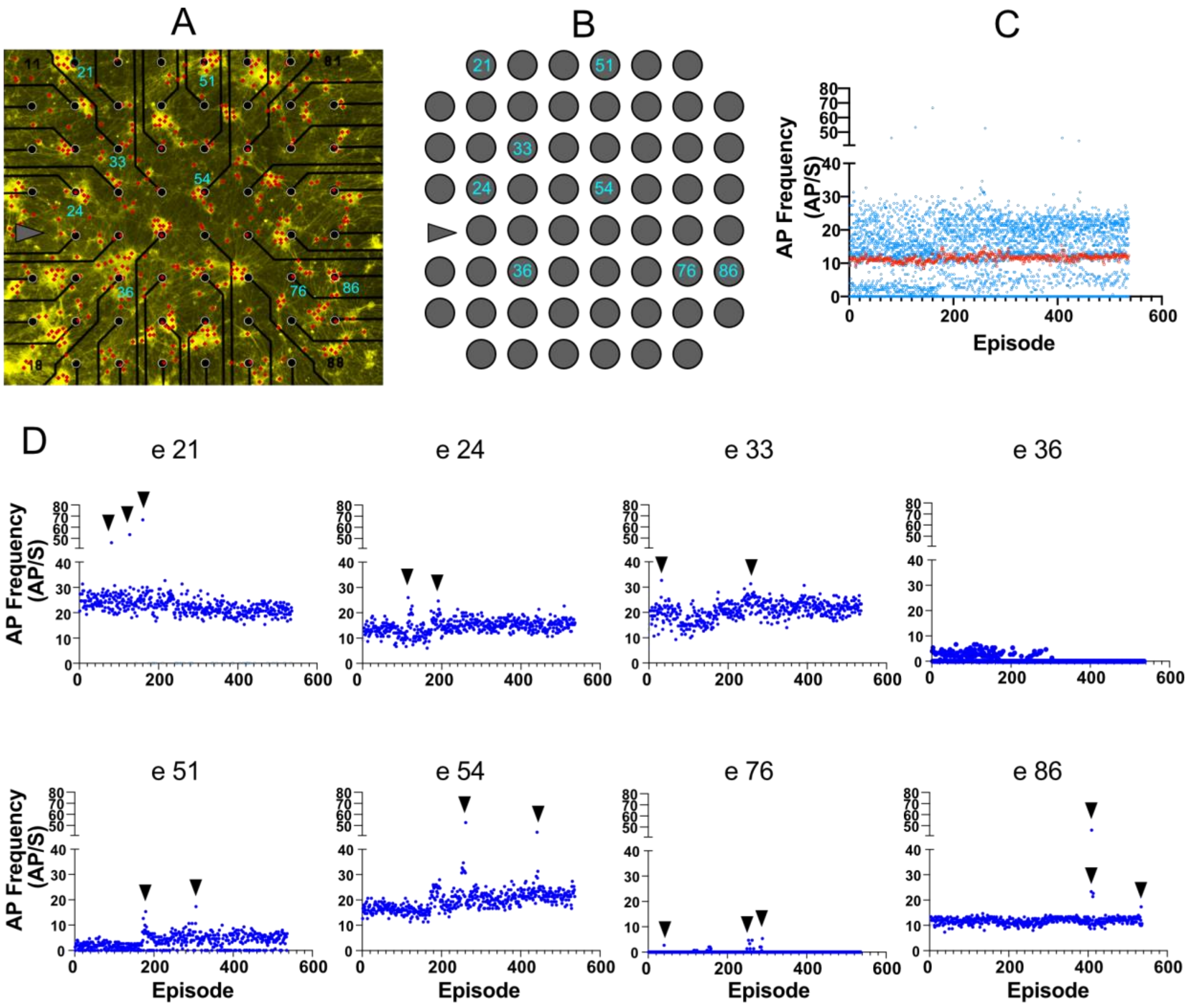


**Figure S7** Tracking the activity during subsequent episodes of holographic stimulation. (A) Network morphology with 536 selected neurons (red dots). (B) Position of the electrodes of interest in the network. (C and D) Average activity in each holographic episode calculated (blue dots) and plotted based on episode number (episode 1 to 536). Each episode included 20 pulses of holographic stimulation applied to the same neuron. Each electrode showed high-amplitude direct responses to a few episodes that targeted particular neurons in the network (indicated by black arrows). Average response of all electrodes (C) did not change during 536 subsequent holographic stimulations (red squares). Each blue dot represents the average response of each electrode to a specific episode.

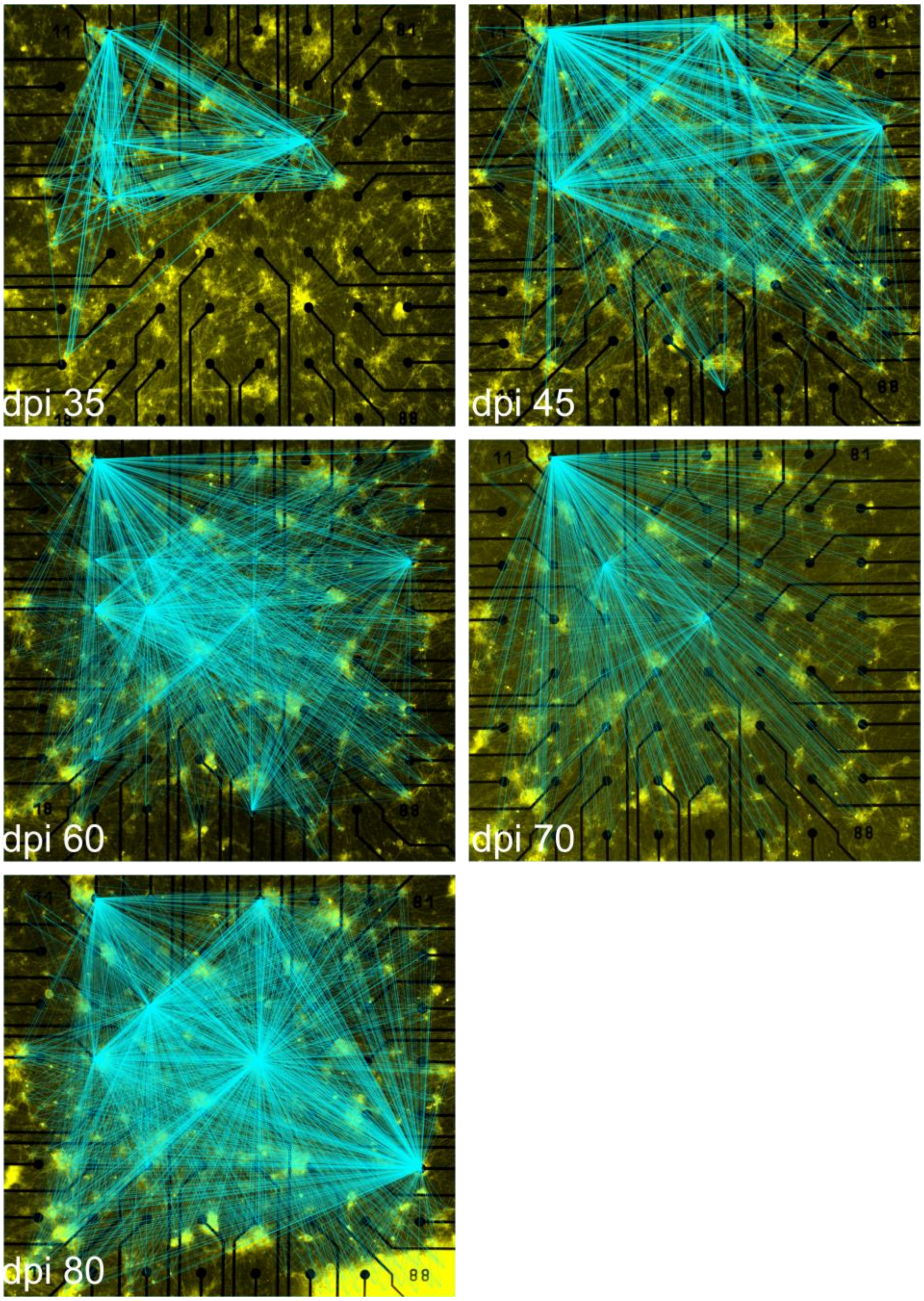


**Figure S8** Network connectivity map at different days post induction overlaid on the fluorescence image of the sample.

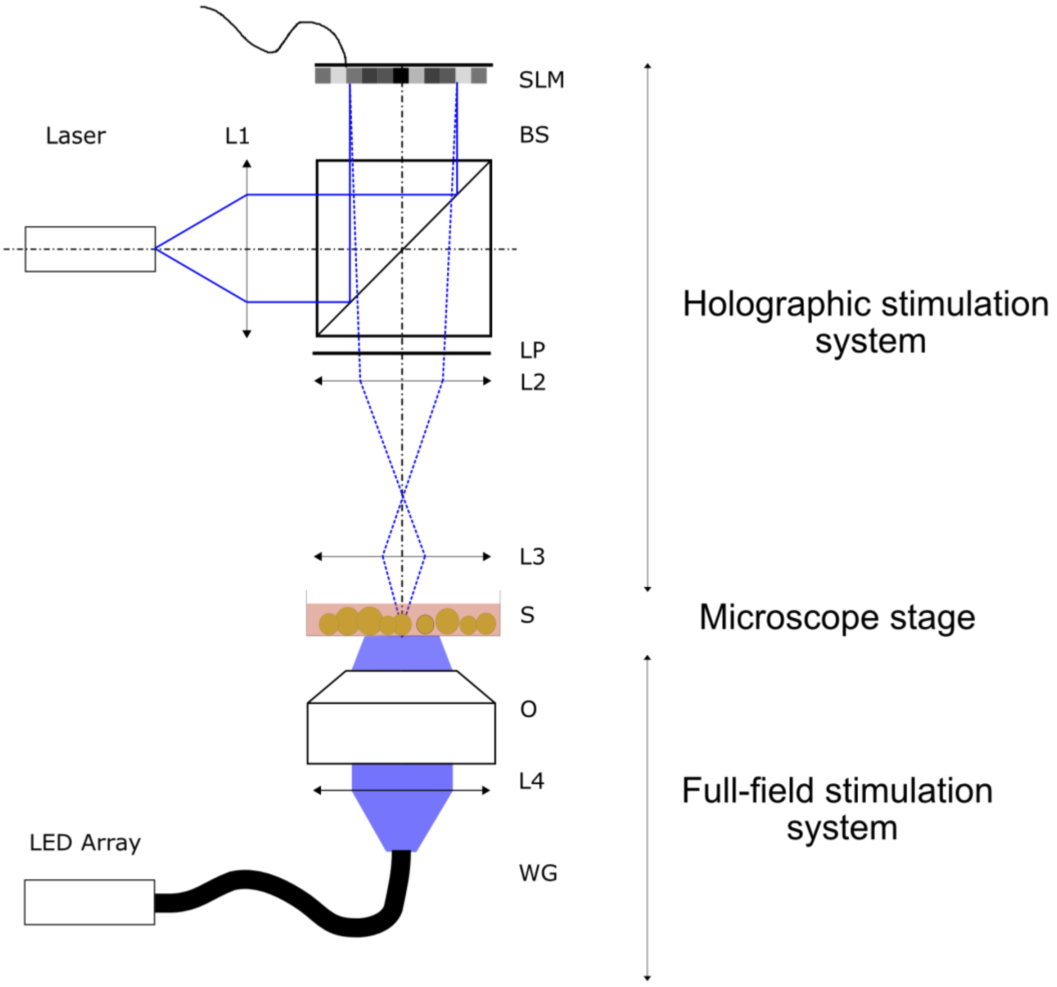


**Figure S9** Optical setups for full-field and holographic stimulations. For full-field stimulation light from an LED array (470 nm) of spectra4 light source is coupled to a liquid waveguide (WG) and outcoupled to an inverted Nikon microscope. Light is directed onto the sample (S) by the tube lens L4 and a microscope objective (O). For holographic stimulation the laser (450 nm laser diode) is collimated by L1 and directed orthogonally onto the SLM by the beam splitter. Modulated light is filtered in polarization to achieve a phase shift of pi and then de-magnified onto the sample by telescope L2-L3. L1 to L3: lenses. SLM: ferroelectric liquid-crystal spatial-light modulator. BS: Non-polarizing 50:50 beam splitter. LP: linear polarizer. S: sample.

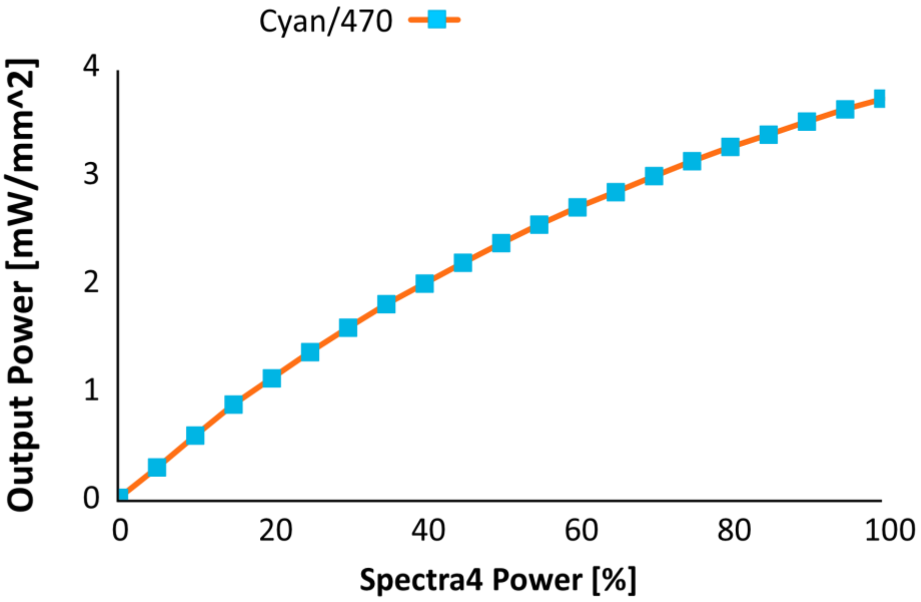


**Figure S10** light intensity of full-field stimulation by spectra4 light source. Light intensity measured on the surface of the MEA chip.
